# Oscillatory-Quality of sleep spindles: from properties to function

**DOI:** 10.1101/2023.06.28.546981

**Authors:** Cristina Blanco-Duque, Suraya A. Bond, Lukas B. Krone, Jean-Phillipe Dufour, Edward C.P. Gillen, Martin C. Kahn, Ross J. Purple, David M. Bannerman, Edward O. Mann, Peter Achermann, Eckehard Olbrich, Vladyslav V. Vyazovskiy

**Affiliations:** Department of Physiology, Anatomy and Genetics, University of Oxford, UK; Sleep and Circadian Neuroscience Institute, University of Oxford, UK; Department of Brain and Cognitive Sciences, Massachusetts Institute of Technology, USA; University Hospital of Psychiatry and Psychotherapy, University of Bern, Switzerland; Centre for Experimental Neurology, University of Bern, Switzerland; Cavendish Laboratory, Astrophysics, University of Cambridge, UK; Department of Experimental Psychology, University of Oxford, UK; Institute of Pharmacology and Toxicology, University of Zurich, Switzerland; Max Planck Institute for Mathematics in the Sciences, Germany

**Author notes:** Corresponding Authors: Cristina Blanco-Duque, Department of Brain and Cognitive Sciences, Massachusetts Institute of Technology, 43 Vassar St, Cambridge, MA 02139, United States, Vladyslav V. Vyazovskiy, Department of Physiology, Anatomy and Genetics, University of Oxford, Sherrington Building, Sherrington Rd, Oxford OX1 3PT, United Kingdom.

**Keywords:** sleep spindles, damping, cortical network synchrony, sleep homeostasis, GRIA1, schizophrenia, sensory stimulation

## Abstract

Sleep spindles are traditionally defined as 10-15Hz thalamo-cortical oscillations typical of NREM sleep. While substantial heterogeneity in the appearance or spatio-temporal dynamics of spindle events is well recognised, the physiological relevance of the underlying fundamental property - the oscillatory strength - has not been studied. Here we introduce a novel metric called *oscillatory Quality* (*o-Quality*), which is derived by fitting an auto-regressive model to short segments of electrophysiological signals, recorded from the cortex in mice, to identify and calculate the damping of spindle oscillations. We find that the *o-Quality* of spindles varies markedly across cortical layers and regions and reflects the level of synchrony within and between cortical networks. Furthermore, the *o-Quality* of spindles varies as a function of sleep-wake history, determines the strength of coupling between spindles and slow waves, and influences the responsiveness to sensory stimulation during sleep. Thus, the *o-Quality* emerges as a metric that, for the first time, directly links the spatio-temporal dynamics of sleep spindles with their functional role.

## Introduction

Brain networks have an intrinsic capacity to generate and sustain a wide range of neural oscillations, which are thought to be a fundamental basis for cognition and behaviour^1–3^. Multi-site recordings of brain activity combined with time-frequency analyses have provided a fundamental understanding of the network dynamics and functional role of brain oscillations^4, 5^. As such, it has long been appreciated that the functional significance of brain oscillations depends on properties such as their topography, frequency, and density^1, 6, 7^. There are, however, fundamental properties of oscillatory systems, such as their damping, which have not been systematically studied.

Damping is a metric frequently used in physics and engineering that measures the decay in the amplitude of an oscillation over time^8^, and therefore reflects levels of oscillatory strength and stability^9^. Damping has recently proven a useful metric for the detection of oscillatory brain activity, like sleep spindles or alpha bursts, which are believed to occur in a form of discrete events^10–14^. Nonetheless, the potential relevance of the oscillatory strength of brain activity for defining network dynamics or functional significance has not been investigated.

This is particularly the case for sleep spindles, which represent one of the most widely studied brain oscillations. Spindles are classically defined as bursts of oscillatory brain activity at frequencies of ∼10-15Hz and durations of 0.5-3 s in rodents^3, 15–21^. These oscillations are thought to play a key role in brain-wide dynamics during sleep, and growing evidence suggests that they may support offline information processing^22–29^ or protect sleep from sensory disruption^30–38^.

Spindles arise in the reticular nucleus of the thalamus^16, 25, 39–41^ and express focally or across widespread thalamo-cortical networks, where they are widely recognized to show high heterogeneity in their frequency, shape, and topography^42–50^. Whether spindles show variability in oscillatory strength, and whether this variability has a physiological relevance for their dynamics and function has, however, been widely disregarded. This may in part be related to the fact that traditional methods to detect spindles typically accept or reject events based on fixed thresholds imposed on parameters such as amplitude or damping, without further assessing whether spindle-to-spindle variability in terms of these metrics may carry important information.

To address this important gap, we developed an approach to quantitatively measure the strength of spindle oscillations and test its relevance at a neurophysiological and functional level by combining multi-site recordings of cortical LFPs and neuronal activity, sleep deprivation, transgenic manipulations, and sensory stimulation in mice. The method is based on a time-frequency analysis, utilising autoregressive modelling of short segments of EEG signals introduced earlier by Olbrich & Achermann^14^ for human EEG. Here we extended this method to allow not only the detection of spindles, but to characterise spindle-events based on their damping. As this metric allows to quantitatively describe the variability of spindles in terms of their oscillatory strength, we called it the oscillatory-Quality (*o-Quality, oQ*).

We undertook comprehensive analyses of spindle *o-Quality* using several lines of enquiry and experimental paradigms. This included estimating the *o-Quality* of spindles across cortical regions and layers, during spontaneous sleep and after sleep deprivation, as well as recordings in wild type and transgenic mice lacking the GluA1 subunit of the AMPA receptor, which were previously found to present deficits in EEG spindles^51^. We further looked at the relationship between sleep spindles and slow waves as a function of their *o-Quality,* and investigated whether the *o-Quality* of spindles is related to sensory responsiveness to auditory stimulation during sleep. Invariably, we find that it is not merely the all-or-none incidence of spindles that matters, but instead their *o-Quality*, which emerges as the key variable reflecting the network dynamics and functional role of spindles.

## Results

### Spindles show substantial variability in their oscillatory strength

We performed continuous electrophysiological recordings of the EEG from frontal, parietal and occipital regions combined with multichannel local field potentials (LFP) from the primary somatosensory (S1, n=21 mice) or primary motor (M1, n=7 mice) cortices (Fig.S1.A-F), in undisturbed freely-moving mice, entrained to a 12:12 light-dark cycle. As expected, all mice slept predominantly during the light phase, of which they spent 84.3 ± 1.28 % of time in NREM sleep (Fig.S1B). Visual inspection of the signals confirmed the occurrence of bursts of oscillatory activity at the spindle frequency (10-15Hz)^6, 52, 53^, in both the EEG and the LFP signals, which showed marked variability in their occurrence and characteristics across time, cortical layers and cortical areas (Fig.1A-B). For example, within a specific location (i.e. specific recording channel), some events were clearly distinct from background activity, while other events were barely discernible. On the other hand, some putative spindle events were prominent across widespread cortical areas (i.e. several LFP channels or different brain regions), while others were readily observed only in one or two channels.

**Fig. 1.**
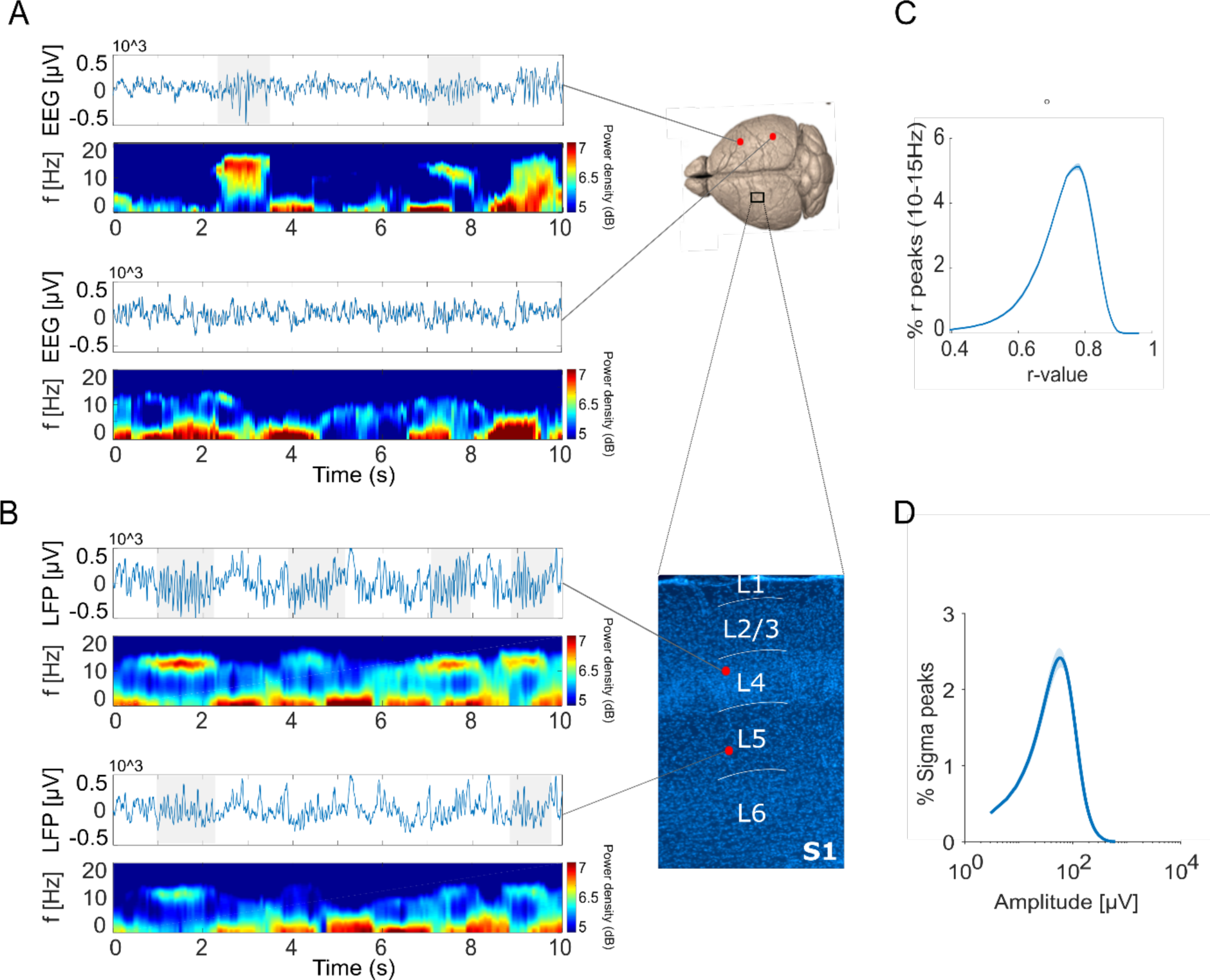
Spindles show a substantial variability in their oscillatory strength. **(A-B)** Ten-second signal segments and respective spectrograms for signals recorded simultaneously from the frontal EEG (A - top) and occipital EEG (A - bottom), layer 4 of S1 (B- top) and layer 5 of S1 (B – bottom). Spectrograms are colour-coded on a logarithmic scale. **(C)** Distribution of the maximum *r* value across poles with frequencies (*f*_*k*_) between 10-15 Hz for an LFP signal recorded from layer 4 in S1. **(D)** Peak sigma (10- 15 Hz) amplitude distribution for the same LFP signal used in panel C (layer 4 in S1). (*Note:* in C and D, line=mean across 7 mice. Shaded area= SEM. EEG: electroencephalogram. LFP: local field potential. S1: primary sensory cortex. SEM: standard error of the mean).

This variability in spindle-like activity is well known to researchers, and was also confirmed in our data set (Fig.1C-D). For example, using auto-regressive modelling and plotting the inverse damping distribution *r* of oscillators (larger r-values correspond to lower damping) with frequency between 10-15Hz during NREM sleep in LFP signals recorded from S1 revealed that oscillatory activity between 10-15 Hz shows a continuous variation in its damping (Fig.1C). This was consistent with the observation of a continuous distribution of LFP amplitudes after band-pass filtering of LFP signals from NREM sleep between ∼10-16 Hz (Fig.1D) - a procedure widely used in the literature to detect spindle events^54–56^.

The key premise for this study was the notion that focusing merely on quantitative measurements of spindle activity (e.g. single amplitude or damping-based detection thresholds to determine incidence) does not take into account the strength of individual spindle events (i.e., how “strong” the oscillatory activity in the spindle-frequency range is during a specific spindle event). This is not merely a methodological issue that can be satisfactorily addressed with the advent of more sophisticated approaches for threshold optimisation. Instead, it highlights the likely possibility that the variability in spindle characteristics has an important meaning beyond what the scrutiny of arbitrarily defined events can provide. We propose that the variability of spindle activity in terms of oscillatory strength represents a fundamentally important dimension that can provide new insights into the underlying neurophysiological mechanisms and function of spindles.

### The oscillatory-Quality: a quantitative metric of spindle activity strength

In the core of our approach is an algorithm that detects oscillatory events in brain signals based on their damping^10, 14^, a measurement used to parameterize oscillatory strength^9^. For the first time, we applied this model to detect spindles on mouse EEG and LFP signals, and, crucially, we extended this approach to allow characterising spindles based on varying levels of damping.

The algorithm consists of fitting an auto-regressive (AR) model of order p=8 to 1-s segments of LFP and EEG signals, shifted by 1 sampling interval throughout the data, which results in a model with a maximum of p/2 oscillators with damping and frequency varying in time (Fig.2A-D). The coefficients of the AR(8)-model are used to calculate an *r*_*k*_ coefficient (with *k* indicating the corresponding modelled oscillator), whose negative logarithm is proportional to the damping constant, therefore *r*_*k*_=1 means no damping and *r*_*k*_=0 means maximum damping (see methods). When the signal is dominated by rhythmic activity like a spindle event (Fig.2A-B), this activity is reflected by a decrease in damping and hence an increase in *r*_*k*_ (Fig.2C), in an oscillator with the corresponding frequency *f*_*k*_ (Fig.2D). Events are detected when the *r*_*k*_ of an oscillator with frequency *f*_*k*_ exceeds a predefined detection threshold (r*_b_*),^10, 14^ and detections are tagged with their respective maximum *r*_*k*_ value and the *f*_*k*_ where *r* is maximum (Fig.2E). The majority of events detected with this approach during NREM sleep were clustered in the traditionally accepted spindle-frequency range in rodents (10-15 Hz) and in the delta range (Fig.S2A). For subsequent analyses, we selected events with tagged *f*_*k*_ between 10-15 Hz.

**Fig. 2.**
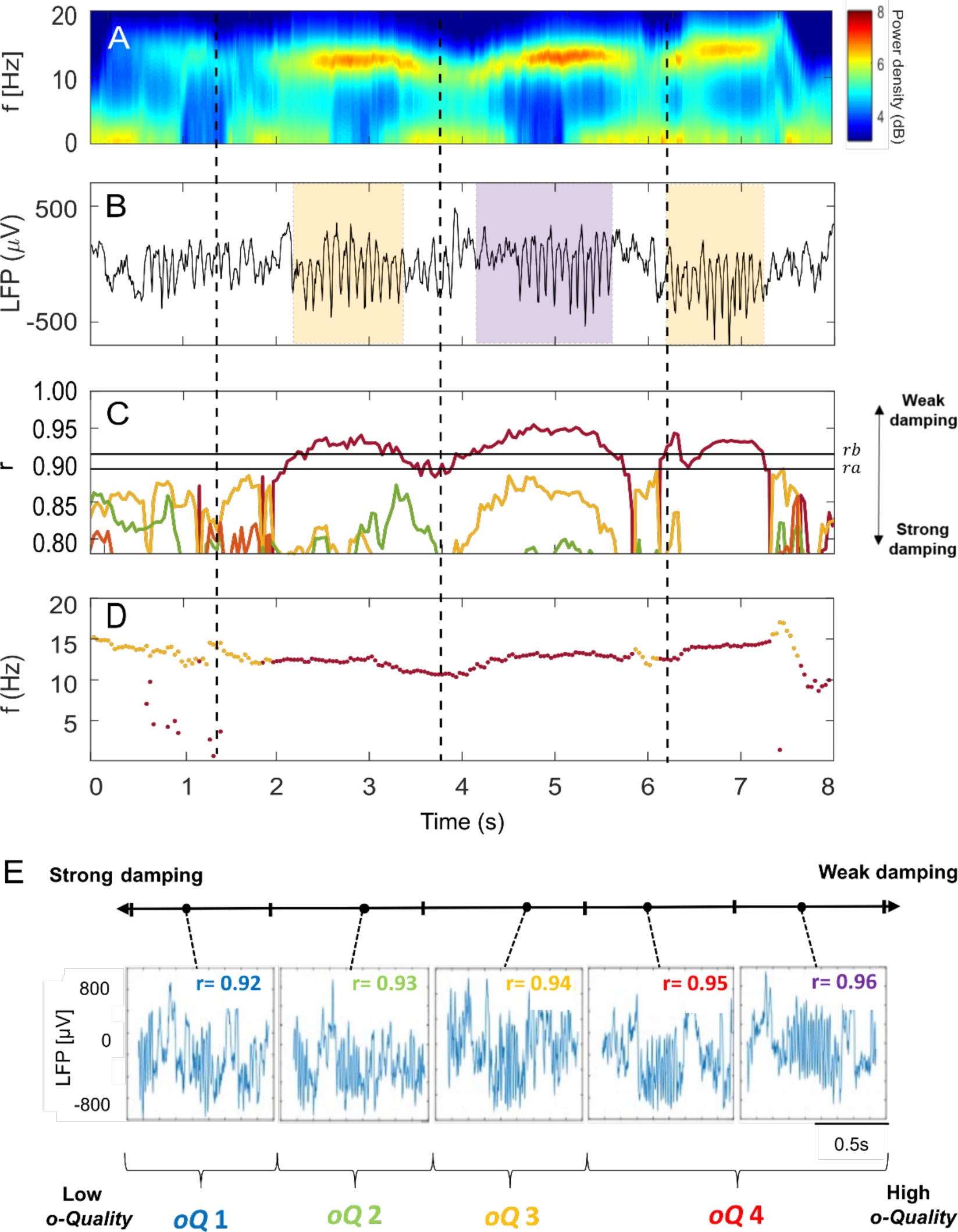
The *Oscillatory-Quality*: a quantitative metric of spindle activity strength. **(A)** Spectrogram of an 8-s segment of LFP recording from S1 in one mouse. Spectra are colour-coded on a logarithmic scale. **(B)** Eight-second segment of LFP data during NREM sleep showing a sequence of detected spindle events, highlighted by shaded coloured boxes. The yellow boxes indicate spindle events whose max-*r* values reached 0.94, while the purple box indicates a spindle event whose max-*r* value reached 0.95. **(C)** Absolute *r* values for the four poles estimated by the AR(8)-model. Each pole is represented with a different colour. The black horizontal lines represent the upper threshold used for detection of oscillatory events (i.e., *r*_*b*_=0.92) and the lower threshold (*r*_*a*_=0.90) used to merge or separate consecutive oscillatory events. **(D)** Frequencies *f*_*k*_ of the poles with lowest damping. **(E)** Examples of spindle events with different levels of damping (i.e. different maximum r values). The maximum *r* value for each detected spindle was used to group spindles into four *o-Quality* groups (oQ1 to oQ4) such that strong-to-weak damping corresponds to low-to-high *o-Quality*. (*Note:* LFP: local field potential. S1: primary sensory cortex. AR: autoregressive).

In engineering and physics, the level of damping in oscillatory systems^8, 9^ is parameterized in terms of a Quality-Factor^8^. In analogy to this metric, we defined an index to parameterize the damping level (i.e. oscillatory strength) in brain oscillations, which we refer to as *oscillatory-Quality* (*o-Quality; oQ*). Specifically, we used the maximum *r* value detected for each event, to group spindles into four *o-Quality* groups (*oQ*1 to *oQ*4) such that strong-to-weak damping corresponds to low-to-high *o-Quality*. These groups were set such that spindles with maximum *r* value between 0.92≤r<0.93, 0.93≤r<0.94, 0.94≤r<0.95 and 0.95≤r would be classified respectively as *oQ*1, *oQ*2, *oQ*3 and *oQ*4 (Fig.2E). Notably, apart from taking into consideration the variability of spindles in their oscillatory strength, this approach does not assume any specific oscillatory waveform and does not require signal filtering in any specific frequency band. This circumvents the potential signal distortion that band-pass filters may generate^57, 58^ and makes this approach suitable for spindle analysis in other animal species and humans with different conditions that may add variability to spindle features^3, 59^.

As expected, we observed that the *o-Quality* of sleep spindles showed a positive relationship with their duration (F_3,18_=674.7, *p*<0.0001) (Fig.S2B), amplitude (measured as the maximum value of the Hilbert transform of the signal during individual events, F_3,18_=62.1, *p*<0.0001) (Fig.S2C), but also frequency (F_3,18_=21.34, *p*<0.001; moderate effect) (Fig.S2D). Spindles with low *o-Quality,* however occurred at a significantly higher rate (F_1.3,7.5_=137.9 GG, *p*<0.0001) (Fig.S2E) than high *o-Quality* spindles. Additionally, spindles with high *o-Quality* showed a higher power in the spindle frequency range than low *o-Quality* spindles (Fig.S3F). Having developed an approach to quantify spindle oscillatory strength, we next explored its utility for furthering our understanding of the spatio-temporal dynamics and function of sleep spindles.

### The spatial dynamics of sleep spindles are reflected in their oscillatory-Quality

First, we posited that if the variability across spindle events in terms of *o-Quality* is biologically meaningful, it should be related to their spatio-temporal dynamics. As previous studies indicate that incidence and frequency of sleep spindles varies as a function of brain region and cortical area^42–50^, we hypothesised that *o-Quality* of spindles will also show topographical gradients. Consistent with this prediction, we found that the incidence of EEG spindles with different *o-Quality* (derivation x *o-Quality* interaction: F_6,54_=4.96, *p<*0.01) (Fig.3A, left), as well as the proportion of high *o-Quality* events detected on EEG signals (F_2,20_=6.45, *p*<0.01) (Fig.3B, top), varied between cortical regions. Generally, across the frontal, parietal and occipital cortex, EEG spindles with a higher *o-Quality* index were more predominant in more anterior cortical areas (Fig.3B, top).

**Fig. 3.**
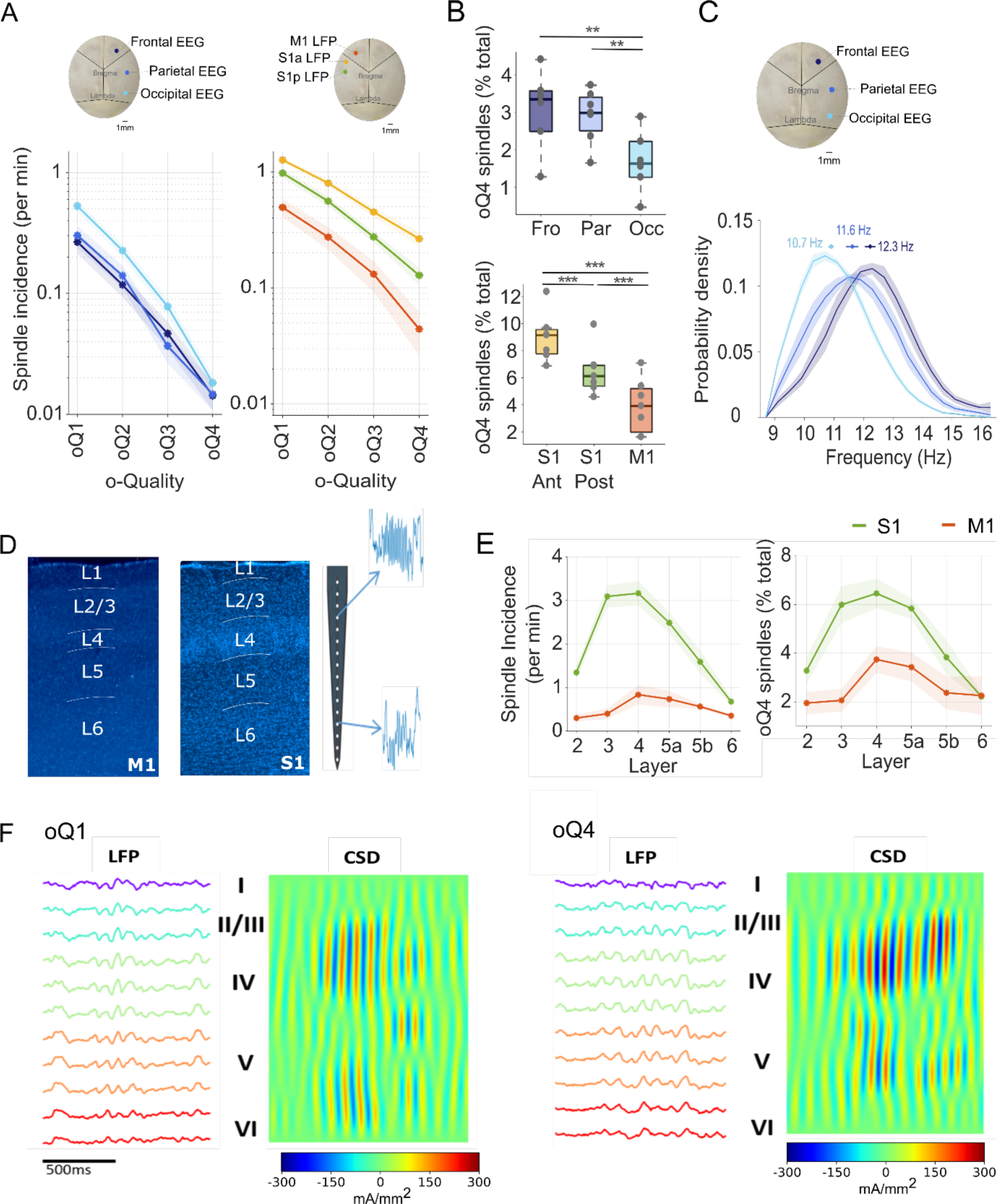
The *o-Quality* reflects spatial dynamics of sleep spindles. **(A)** Incidence per minute of spindles detected in EEG (frontal, parietal and occipital) and LFP (anterior S1, posterior S1 and M1) derivations as a function of spindle *o-Quality*. Dots= mean across mice; shadows=SEM. **(B)** Number of high *o-Quality* (o-Quality 4) spindles as a percent of total spindles detected in EEG (frontal, parietal and occipital) and LFP (anterior S1, posterior S1 and M1) derivations. *For boxplots*: black lines= mean across mice, boxes= SEM, whiskers= 95% confidence intervals, dots= individual values for each mouse. *p<0.05, **p<0.01, ***p<0.001. **(C**) Frequency (Hz) distribution for spindles detected in different EEG derivations (frontal, parietal and occipital). Lines=mean across mice; shadows=SEM. **(D)** Histological verification of probe location across cortical layers in M1 and S1. Illustrations showing examples of spindle events detected in cortical layers 4 and 6 of S1 (right). **(E)** Mean spindle incidence per minute (left) and percentage of detected high o-Quality (oQ4) spindles (percentage of total number of spindles; right) across different cortical layers of S1 and M1 cortices. **(F)** Example *o-Quality* 1 and *o-Quality* 4 spindles with LFP and current source density (CSD; red: current source, blue: current sink) signal of primary somatosensory cortex. Layer centroids are marked by roman numerals. (*Note:* EEG: electroencephalogram. LFP: local field potential. S1: primary sensory cortex. M1: primary motor cortex. SEM: standard error of the mean. CSD: current source density).

Consistent with the finding of a positive relationship between intra-spindle frequency and *o-Quality* (Fig.S2D), we observed that the predominant frequency of EEG spindle events varied among cortical areas, with slowest spindles occurring in the occipital cortex (F_2,20=_16.38*, p*<0.001) (Fig.3C). These results are in line with previous mouse EEG studies^44, 50^. Conversely, human studies have reported that spindles show an anteroposterior increase in their frequency^48, 60–65^.

Likewise, the distribution of LFP spindles as a function of their *o-Quality* (*oQ*1 - *oQ*4) varied between cortical regions (derivation x *o-Quality* interaction F_2.56,54_=19.81, *GG*, *p*<0.01) (Fig.3A, right). Specifically, we found that spindles recorded with LFP electrodes from two locations within S1, were of higher *o-Quality* in more anterior locations and overall showed higher *o-Quality* than spindles recorded with LFP electrodes from M1 (F_2,20_=13.38, *p*<0.001) (Fig.3A, right - 3B, bottom). These results suggest that the oscillatory strength of spindles is not homogenous across the cortex, but shows distinct topographic gradients, consistent with established morphological and functional differences between cortical areas.

In contrast to the variability of spindles across cortical regions^43–45, 48–50, 62, 63^, their laminar dynamics have received much less attention^66–69^. To the best of our knowledge, layer-specific changes in damping of spindle oscillations has not been previously studied. To this end, we compared the incidence and *o-Quality* of spindles recorded during NREM sleep along 16-channel laminar probes implanted in S1 and M1 (Fig.3D). We found that in both S1 and M1, the total incidence of spindles, and high *o-Quality* events in particular, were highly layer- and region-specific (Fig.3E). The most prominent spindle activity (F_5,30_=30.2, *p*<0.0001), and of the highest *o-Quality* (F_5,30_=10.3, *p*<0.0001), was found between layers L2/3-deep, L4 and superficial subdivision of L5. This distribution shifted to somewhat deeper electrodes in M1, where the incidence (F_1.9,11.4_=4.03, *GG*, *p*<0.05) and the proportion of high *o-Quality* spindles (M1 F_5,30_=3.65, *p*<0.05) was higher in L4 and superficial subdivision of L5 (Fig.3E). This result is consistent with anatomical evidence indicating that thalamo-cortical projections to M1 form most synapses in L5 and L4, and to a lesser extent in L2 and L3^70^. Thus, our data suggest that not only spindle incidence but prominently their *o-Quality* vary as a function of both cortical area and cortical layer.

The question arises whether spindles with different *o-Quality* may have different generators. To address this, we compared the LFP and current source density (CSD) magnitude in different layers during LFP spindles with different *o-Quality* detected in S1 (Fig.3F). LFP or CSD magnitudes were calculated as the 1-second root-mean-square (RMS) value centred around each spindle’s maximum-envelope LFP cycle. Across mice, the average spindle laminar profile was consistent, with maximal LFP and CSD amplitudes in layers 2/3 and 4, as described above. The LFP signal amplitude then decreased in layer 5 and yet further in layer 6 (Fig.S3). A second, smaller CSD signal was observed in the deeper channels in every animal (Fig.3F). A repeated-measures ANOVA on the laminar LFP and CSD root mean square (RMS) values revealed significant effects of layer and *o-Quality* on the signal magnitude (*p*<0.001 in each mouse, LFP and CSD). The laminar profile of CSD was, however, only weakly affected by the spindle *o-Quality* (i.e. significant interactions between layer and *o-Quality* (*p*<0.001) but only small effect sizes (partial eta-squared < 0.1).

Using the layer magnitudes of all unique spindle events, we performed a principal component analysis (PCA) for every mouse separately. LFP and CSD amplitudes across layers were highly correlated: the first principal component in every mouse was the only component with an eigenvector above 1, with explained variances ranging from 71% - 84% for the LFP and 71% - 81% for the CSD. No distinct clusters were discernible in any mouse for either LFP or CSD, further suggesting that one fundamental laminar profile is indeed highly dominant across all spindles, and spindles with different *o-Quality* have similar generating networks.

### The degree of network synchrony underpins the oscillatory-Quality of sleep spindles

The observation that *o-Quality* of spindles correlated with both their duration (Fig.S2B) and amplitude (Fig.S2C), suggested that this metric may reflect the size of the network involved, or the degree of network synchronisation during spindling. It is well-known both from human and animal studies, that spindles can occur in widespread cortical areas, but most spindles are expressed in restricted local areas^43, 49, 50, 67, 68^. In line with this, visual inspection of LFP and MUA signals, recorded with multi-channel probes that spanned either vertically across cortical layers or horizontally across cortical areas (Fig.4A), revealed that spindles in S1 and M1 display a significant diversity in their spatial extent (Fig.4B). In some cases, spindles occurred at the same time in most recording sites, including both the LFP and EEG. In other cases, sometimes just a few seconds later, only a few channels manifested discernible spindling at a given time.

**Fig. 4.**
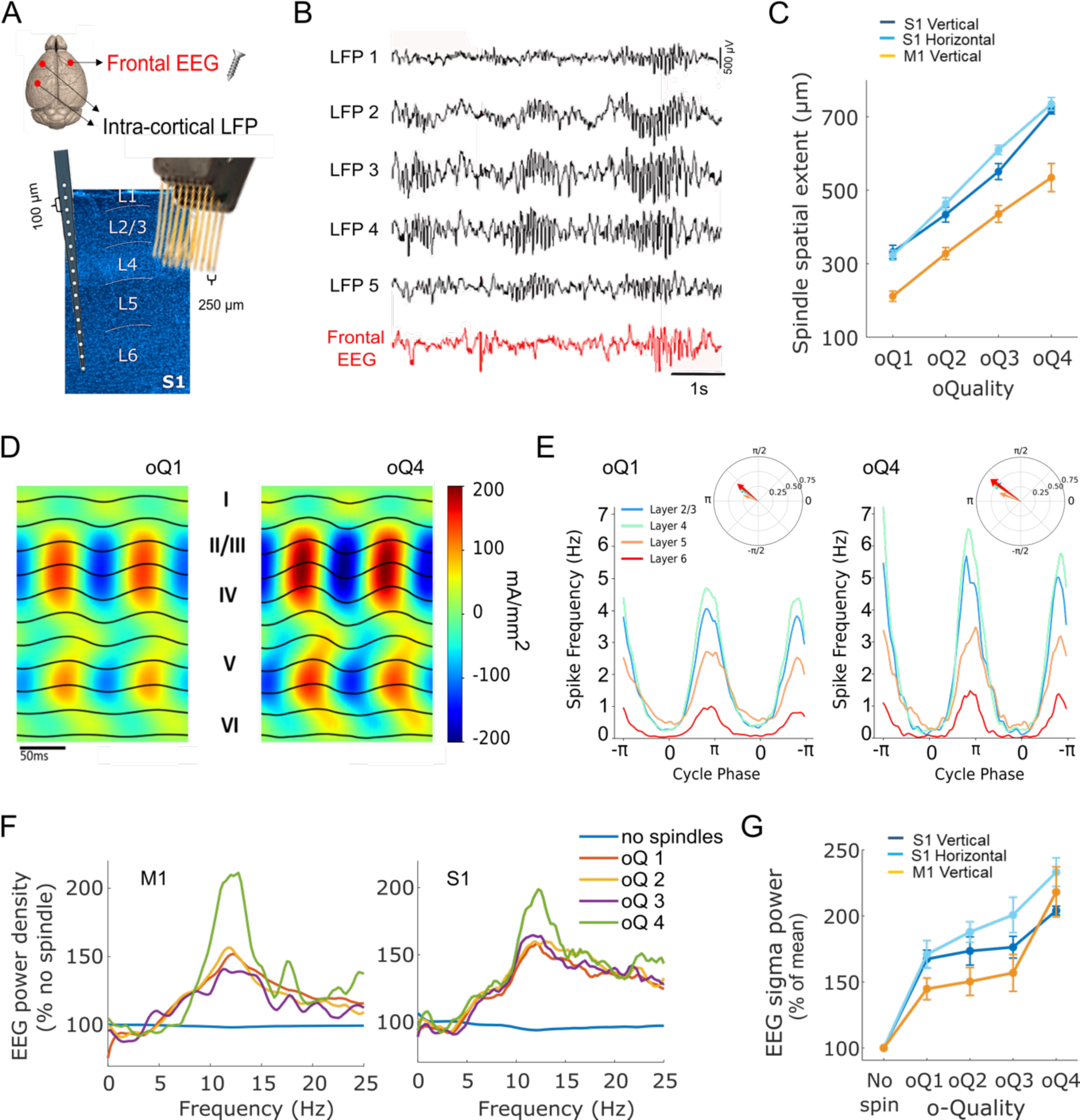
The spindle o-Quality reflects synchrony within local and global cortical networks. **(A)** Diagram indicating the location where a frontal EEG screw (n=21 mice) and intracortical probes (S1 and M1) were implanted. All mice (n=21) were implanted with a frontal screw. Additionally, a S1 microwire array was implanted in n=7, a S1 laminar probe in n=7 and a M1 laminar probe in n=7 of these mice. **(B)** Representative LFP signals (6 s of NREM sleep in one mouse) recorded from five contiguous electrodes within a laminar probe implanted in S1 (black) and corresponding frontal EEG signal (red). These traces show examples of local spindles (restricted to a few LFP channels) and a more global spindle that expresses in all the LFP channels and the EEG. **(C)** Mean spatial extent of LFP spindles recorded from both laminar probes (in S1–dark blue and M1-orange) and micro-wire arrays (S1–light blue), as a function of spindle *o-Quality*. The spatial extent of a spindle event is calculated from the number of LFP electrodes involved in that specific event. **(D)** Mean maximum-envelope 2-cycle average in an example mouse, for qualities 1 and 4. LFP traces are superimposed on the spatially smoothed CSD, averaged across all spindles in that mouse. Left and right plots share the same y-axis and colour range. **(E)** Multi-unit activity by layer. Cycle phase is obtained from the Hilbert transform of the LFP in a layer 4 channel. Compass plot shows mean firing angle and resultant vector length for all spikes within 1 s of every spindle centre for each layer. Rayleigh’s test of circular uniformity confirmed significant phase coupling between MUA and LFP phase in every layer, *oQuality* and mouse. **(F)** Mean EEG power density in the frontal EEG channel during epochs with detected LFP spindle events with different *o-Quality* or during spindles without detected LFP spindles (blue) recorded in M1 (left) and S1 (right). Mean values are shown for each frequency bin expressed as a percentage of mean power density during NREM sleep epochs without detected spindles (‘% no spindle’). **(G)** Mean EEG sigma power in the frontal derivation during epochs with detected spindles as a function of the *o-Quality* of LFP spindles detected in S1 (dark blue= laminar probe; light blue= micro-wire array) and M1 (orange). Note: figures show mean and, where relevant, SEM across mice. (*Note:* EEG: electroencephalogram. S1: primary sensory cortex. M1: primary motor cortex. LFP: local field potential. SEM: standard error of the mean. CSD: current source density. MUA: multi-unit activity. S1 laminar: n=7; S1 micro-array: n=7; M1 laminar= S1 laminar: n=7.).

Consistent with our hypothesis, we observed a strong positive association between the spatial extent of LFP spindles and their *o-Quality* in all cortical regions (S1_vertical:_ F_3,18_=196.89, *p<*0.001; M1: F_1.2,7.2_=70.04 *GG*, *p<*0.001; S1_horizontal_: F_1.1,5.4_=327.5 *GG*, *p<*0.001), especially in the S1 area for spindles recorded both within and across cortical layers (*o-Quality* x derivation interaction: F_2.5,21.73_=2.46 *GG*, *p<*0.05) (Fig.4C). In other words, those events of highest *o-Quality* were present simultaneously across the largest number of channels, while events of lowest *o-Quality* were typically restricted to a few recording channels only. In every mouse, around 46% ± 6% (mean ± SEM) of spindles were detected in only one layer and co-occurrence with other layers was a function of layer distance (Fig.S4A). While the laminar profile of spindle detections did not change with increasing *o-Quality* metric, a higher co-occurrence rate was significantly linked to a higher *o-Quality* metric (One-way ANOVA: *p*<0.001, Fig.S4B-C). On average, spindles with lowest *o-Quality* were expressed within a radius ∼280 ± 15.4μm in S1 and ∼150 ± 20.6μm in M1 (i.e. expressed in ∼22% to 50 % of all LFP channels). As the *o-Quality* of spindles increased, the spatial extent of their expression gradually increased, until this reached a radius of ∼680 ± 25.9μm in S1 and ∼515 ± 35.1μm in M1. These results suggest that the *o-Quality* of LFP spindles reflects network synchrony.

Although the precise site of origin of individual spindle events is difficult to determine with our (or indeed any) recording approach, we established that the occurrence of spindles in the LFP signals invariably correlated with MUA modulation in the same recording channels (Fig.4D-E). Invariably, MUA in all layers was significantly coupled to LFP phase (Rayleigh’s test of circular uniformity: p<0.001 in all layers, *o-Quality* groups, and mice), and phase-coupled spiking was most prominent in layers 2/3 and 4 in all animals (Fig.4E). In order to test the relationship between spindle *o-Quality* and spiking activity, the mean firing angle and resultant vector length of one LFP channel per layer were averaged within each mouse, and a two-way repeated-measures ANOVA was performed on the pooled averages. This revealed that the mean firing angle was not significantly affected by either layer (*p* = 0.163) or spindle *o-Quality* (*p* = 0.480), while resultant vector length increased significantly with *o-Quality* (*p*<0.001), but not layer (*p* = 0.238). No significant layer x *o-Quality* interactions were observed (mean firing angle: *p* = 0.635, resultant vector length: *p* = 0.578), suggesting that the spiking pattern remains largely unaffected across *o-Quality* levels, save for higher LFP phase-spiking coupling with higher spindle *o-Quality*. This suggests that the spatial extent of LFP spindle events reflects predominantly locally originating network activity, rather than volume conducted signals, and the *o-Quality* is a reliable measure of how strongly spiking is modulated during spindle oscillations.

An important question arises as to what extent spindle *o-Quality* also reflects synchrony within wider cortical networks. To address this question, we made use of simultaneous recordings of both the LFP and the EEG at distant locations (Fig.4A). First, we assessed the relationship between the occurrence of LFP spindles and the probability of spindling in the distant EEG signal. Consistent with the notion that the majority of spindles are of low *o-Quality,* we found that 91.7% ± 1.3% (mean ± SEM) of all S1 LFP spindles are not accompanied with EEG spindles. Furthermore, the occurrence of low *o-Quality* spindle events in the LFP was associated with a relatively modest increase of EEG spectral power at the spindle frequency range (10-15Hz) during the corresponding epoch, while high *o-Quality* LFP spindle events correlated with a prominent spindle-frequency peak on the corresponding EEG spectra (Fig.4F-G) (main effect of *o-Quality* F_1.48,23.74_=21.34, *GG*, *p*<0.0001, and significant positive quadratic effect of *o-Quality* F_1,20_=5.05, *p*<0.05, on the EEG power density at 12.5Hz). Together these results suggest that the spindle *o-Quality* reflects the synchrony of wide cortical networks involved in the expression of spindle events.

### Spindle o-Quality correlates with the probability of spindle and slow wave coupling

Our data suggest that spindle *o-Quality* varies not only between individual events, but also between cortical regions and layers, and correlates with other spindle characteristics, such as their spatial synchronisation and their amplitude. This raises the possibility that the spindle *o-Quality* metric reflects, more generally, the state of the thalamo-cortical network, which changes dynamically as a function of incoming inputs, the state of arousal or preceding sleep-wake history. Notably, another major sleep oscillation – the slow wave – is also characterised by the occurrence of local and global events, which can travel across the cortex, vary greatly in terms of their amplitude, topography, and spatial extent, and are exquisitely sensitive to the preceding duration of wakefulness and sleep, as well as network excitability^49, 71–81^. To our knowledge, these well-known properties of sleep slow waves have not been directly linked to the oscillatory strength of spindles.

First, we hypothesised that spindle *o-Quality* is directly related to the probability of coupling between individual slow waves and spindle events. This may be the case given that the spatiotemporal synchrony of spindles is driven by cortico-thalamic inputs, in which slow waves play an important role^66, 82–87^, and that our data suggest that spindle *o-Quality* reflects local and global network synchrony. To address this hypothesis, we detected individual depth-positive high-amplitude slow waves (0.5-4 Hz, see Methods) in the EEG and LFP signals recorded from layer 5 of S1 using a previously published algorithm^81^, and determined the probability of an occurrence of spindle events immediately after a slow-wave detection (Fig.5A).

**Fig. 5.**
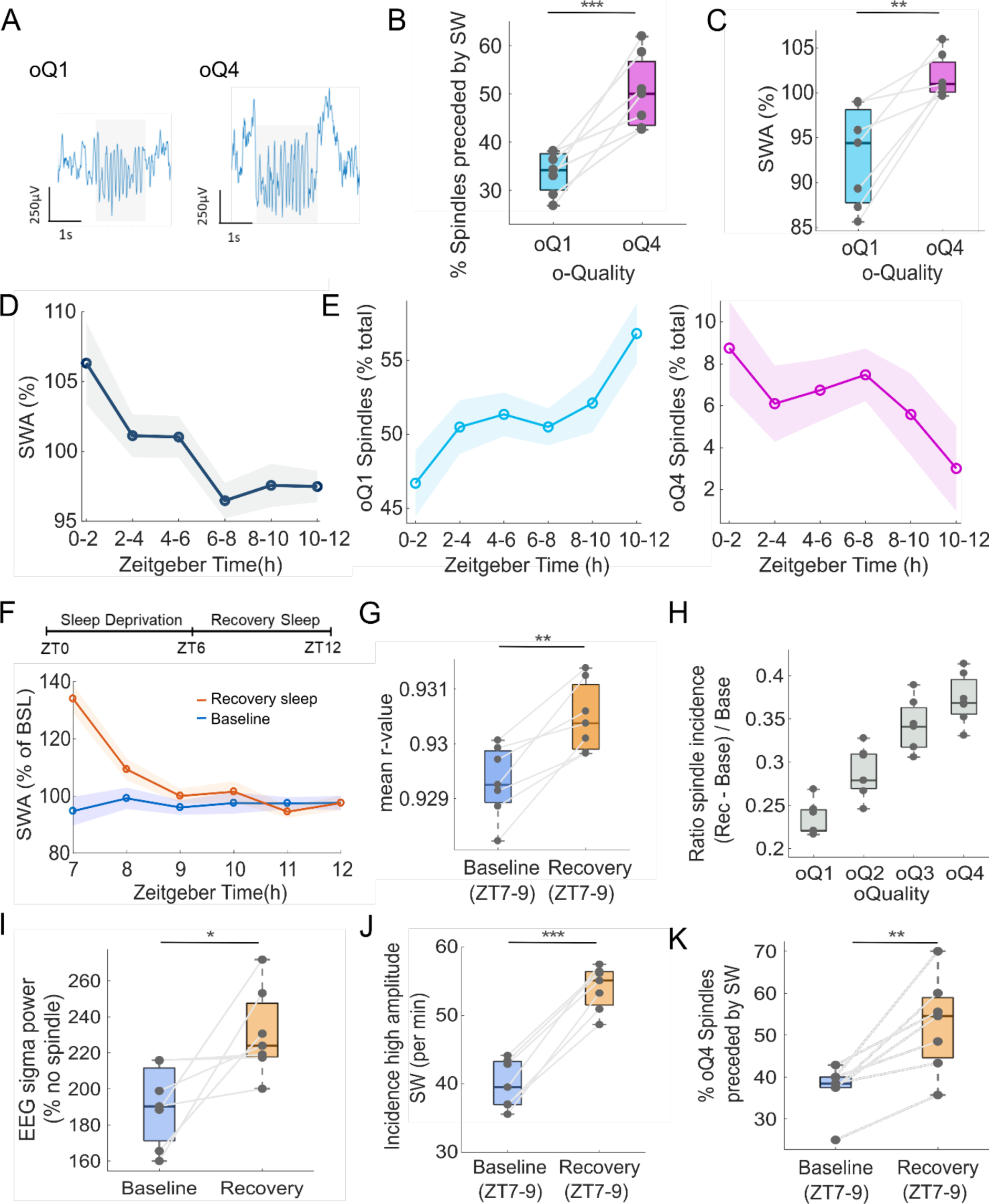
Spindle o-Quality, slow waves and sleep homeostasis. **(A)** Representative examples of spindle events with high (oQ4) and lower (oQ1) *o-Quality* detected in S1. **(B)** Percent of oQ1 and oQ4 spindles preceded by SW. **(C**) LFP power in SWA frequency range (0.5-4 Hz) during 4-s epochs with detected spindles with low (oQ1) and high (oQ4) *o-Quality* values. SWA values are expressed as % of mean 12-h NREM sleep value. **(D)** Time-course of SWA in the S1 LFP signal across a 12-h light period. SWA values are expressed as % of mean 12-h NREM sleep value. **(E)** Incidence of low o-Quality (oQ1) spindles (left) and high o-Quality (oQ4) spindles as a percent of total detected spindles across the 12-h light period (ZT 0-12). **(F)** Time-course of SWA in the S1 LFP signal during sleep recovery after 6 hours of sleep deprivation (orange) and corresponding baseline sleep time-period (blue). SWA values are expressed as % of mean 6-h (ZT7-ZT11) baseline value. **(G)** Mean spindle max-*r* value measured during the first two hours of recovery sleep after sleep deprivation (ZT7-ZT9) and corresponding baseline sleep time-period. **(H)** Spindle incidence ratio between recovery sleep after sleep deprivation and baseline sleep as a function of spindle *o-Quality*. **(I)** EEG power in the spindle frequency range (10-15Hz) during 4-s epochs where oQ4 LFP spindles occurred during baseline (ZT7-9) and recovery (ZT7-9) sleep. The mean EEG sigma power is expressed as a percentage of mean power density during NREM sleep epochs without detected spindles (‘% no spindle’). **(J)** Incidence of high-amplitude SW during the first two hours after sleep deprivation (ZT7-9) and the corresponding baseline sleep time-period. **(K)** Percent of high *o-Quality* (oQ4) spindles preceded by SW during the first two hours after sleep deprivation (ZT7-9) and the corresponding baseline sleep time-period. (*Note:* LFP: local field potential. SWA: slow wave activity. ZT: zeitgeber time. SW: slow waves. For figures D-E, dots=mean across mice; shaded areas=SEM. For boxplots: black lines= mean across mice, boxes= SEM, whiskers= 95% confidence intervals, dots= individual values for each mouse. Analyses were performed on one LFP (S1) channel per mouse, i.e. the channel that showed highest spindle density. *p<0.05, **p<0.01, ***p<0.001.).

In general, we found that 3.42% ± 0.4 of all slow waves were followed by a spindle event within 125ms, and 41.5% ± 2.57% of spindle-events were preceded by a locally recorded slow wave, consistent with the notion that only a subset of spindles are nested in slow waves during physiological NREM sleep^49, 87^. Interestingly, we found a significant positive association between the probability of slow wave and spindle coupling and corresponding spindle *o-Quality*. Specifically, spindle events of higher *o-Quality* (*oQ*4) were, in all individual mice, more likely to be preceded by local slow-waves than low *o-Quality* spindles (*oQ*1) (F_1,6_=31.2, *p*<0.001) (Fig.5B), and LFP power density in the slow wave frequency range was enhanced during 4-s epochs with high *o-Quality* spindle events (F_1,6_=21.61, *p*<0.01; Fig.5C). Notably, the coupling between slow waves and both low *o-Quality* (F_1,6_=199.01, *p*<0.0001) and high *o-Quality* spindles (F_1,6_=162.6, *p*<0.0001), was substantially reduced when the timestamps of slow wave occurrence were shifted offline by 700ms (Fig.S5A-B), which indicates that the slow wave and spindle coupling does not arise by chance.

### Spindle o-Quality reflects network synchrony under increased sleep pressure

Since slow-wave activity (SWA), as well as the incidence of high-amplitude slow waves, is sensitive to preceding sleep-wake history^79–81, 88–92^, we next hypothesised that spindle *o-Quality* may also reflect homeostatic sleep pressure. Both human and rodent studies have suggested an inverse correlation between EEG SWA and spindle activity dynamics^48, 88, 93–95^; however, this relationship varies depending on cortical region, specific properties of slow-waves and spindles, as well as the temporal scale used^46, 47, 51, 93, 96, 97^. We should point out that little is known about the effects of sleep-wake history on the relationship between SWA and spindles in the somatosensory cortex of mice, and how sleep deprivation affects oscillatory strength of spindle activity has not been investigated.

Consistent with previous studies, we found that LFP SWA shows a declining time course across the light period (factor time, F_5,30_=3.63, *p*<0.01, Fig.5D), which is a habitual sleep phase in laboratory mice^47, 98^. However, the time course of sleep spindles across the day varied depending on their *o-Quality* (interaction *o-Quality* x time: F_5,60_=7.77, *p*<0.0001, Fig.5E). Specifically, the incidence of low *o-Quality* (*oQ*1) spindles increased across the 12-h light period (ZT 0-12), (F_5,30_=3.57, *p<*0.01; linear effect F_1,6_=7.35, *p*<0.05), while the incidence of high *o-Quality* (*oQ*4) spindles showed a decreasing time-course across this same period (F_5,30_=8.34, *p<*0.0001; linear effect F_1,6_=39.70, *p*<0.001). From a methodological point of view, this observation suggests that the choice of threshold will have an important impact on the incidence of spindles if their oscillatory strength is not taken into account.

To further address the effects of preceding sleep-wake history on spindle *o-Quality*, next we performed 6-h sleep deprivation (SD), which is a conventional approach to physiologically increase the levels of homeostatic sleep pressure. As expected, LFP SWA increased significantly after SD, which was followed by its gradual decline (interaction condition x time interval F_5,60_=34.40, *p*<0.0001, Fig.5F). Interestingly, early NREM sleep immediately after sleep deprivation was also characterised by an increase in mean *o-Quality* of sleep spindles relative to baseline sleep (F_1,6_=27.67, *p*<0.001; Fig.5G; increased r-value = higher *o-Quality*). Consistently, we also obtained a significant interaction between *o-Quality* (*oQ*1 - *oQ*4) and condition (baseline, recovery) on the incidence of spindles (F_3,36_=6.31*, p*<0.001), suggesting that the effects of sleep deprivation on spindles varied as a function of their *o-Quality.* This conclusion was supported by the observation of a positive relationship between *o-Quality* of sleep spindles and the magnitude of their increase after sleep deprivation (F_3,18_=37.83, *p*<0.0001) (Fig.5H). Additionally, we also found a significant interaction between the *o-Quality* of LFP spindles and sleep condition (baseline, recovery) on the EEG sigma power in the distant EEG signal (F_1,6_=7.28, *p*<0.05, Fig.5I). Specifically, high *o-Quality* LFP spindle events resulted in a prominent spindle-frequency peak on the corresponding EEG spectra, which was significantly higher during the first two hours after sleep deprivation (ZT7-9) compared to baseline sleep (F_1,6_=7.8, *p*<0.05).

Finally, we investigated whether the coupling between slow waves and spindles is affected by preceding sleep-wake history. Consistent with earlier studies^47, 93, 94, 97^, we found an increased incidence of high-amplitude slow waves during the first two hours after sleep deprivation (ZT7-9) compared to baseline sleep (F_1,6_=195.7, p<0.0001, Fig.5J). At the same time, the proportion of high *o-Quality* spindles linked with slow waves was 15.2% ± 3.3% higher during the first two hours (ZT7-9) of recovery sleep after SD compared to the low sleep pressure condition (ZT7-9) during baseline (F_1,6_=21.35, p<0.005, Fig.5K). Notably, this increase in coupling between slow waves and spindles as a function of condition (baseline ZT7-9 vs recovery ZT7-9) was attenuated for low *o-Quality* spindles (F_1,6_=4.54, p=0.09, Fig.S5C), and completely abolished when the timestamps of slow wave occurrence were shifted offline by 700ms (F_1,6_=1.68, p=0.24, Fig.S5D). This indicates that the increase in slow wave and spindle coupling during the first two hours after sleep deprivation does not arise merely by chance but may reflect increased synchronisation of the thalamo-cortical network.

Taken together, these results suggest that spindle *o-Quality* is a metric that is sensitive to the levels of homeostatic sleep need and reflects the state of the thalamo-cortical network under increased sleep pressure. Furthermore, our data provide novel insights into the relationship between the two major sleep oscillations, which we surmise is shaped by the level of network synchronisation.

### GluA1-mediated neurotransmission is essential for the large-scale network synchronization of spindles

Our data demonstrate that spindle *o-Quality* is an informative metric for understanding spatio-temporal synchrony of sleep oscillations. However, the underlying mechanisms linking the network states with oscillatory dynamics remain unclear. To begin addressing the role of spindle *o-Quality* from a mechanistic angle, we next turned our attention to a recently established mouse model of deficient EEG spindle-activity^51^. These animals, which lack the GluA1 subunit of the AMPA receptor and show impaired synaptic plasticity^99–101^, show marked and selective attenuation of spindle power in the frontal EEG during NREM sleep ^51^. The GluA1 subunit plays a key role in a broad range of synaptic functions, including synaptic plasticity^102, 103^. Therefore, this mouse model is a promising tool to investigate network mechanisms of local and global spindle propagation. An additional rationale for choosing this model was that recent genome wide association studies have linked the GRIA1 gene, which encodes GluA1, with schizophrenia^104–106^, and in line with this, GRIA1^-/-^ mice show phenotypes relevant for schizophrenia^51, 107–112^. It is well known that EEG spindle activity is markedly reduced in patients with schizophrenia^113–123^ and therefore spindle *o-Quality* can potentially have a promising and yet untapped clinical relevance in this regard.

We performed chronic EEG and LFP recordings in freely-moving GRIA1^-/-^ mice (n=7) and their wild-type (WT) littermates (n=7), and applied our spindle detection algorithm as described above to the frontal EEG and the LFP recorded in S1. We confirmed ^51^ a marked decrease of EEG spectral power in the spindle frequency range during NREM sleep in GRIA1^-/-^ relative to WT mice (F_80,960_=8.97, *p*<0.0001) (Fig.6A), which, as expected, was associated with a decrease in the total spindle incidence (t_7.9_=5.21, *p*<0.001) (Fig.6B). Further, we also observed that the remaining spindles that were still detectable in the EEG of GRIA1^-/-^ mice, were of significantly lower *o-Quality* than in WTs (t_12_=4.27, *p*<0.001) (Fig.6C; lower r-value = lower *o-Quality*), and were associated with an attenuated increase of EEG power in the spindle-frequency range (Fig.6D-E; Genotype x frequency: F_80,960_=6.47, *p*<0.0001).

**Fig. 6.**
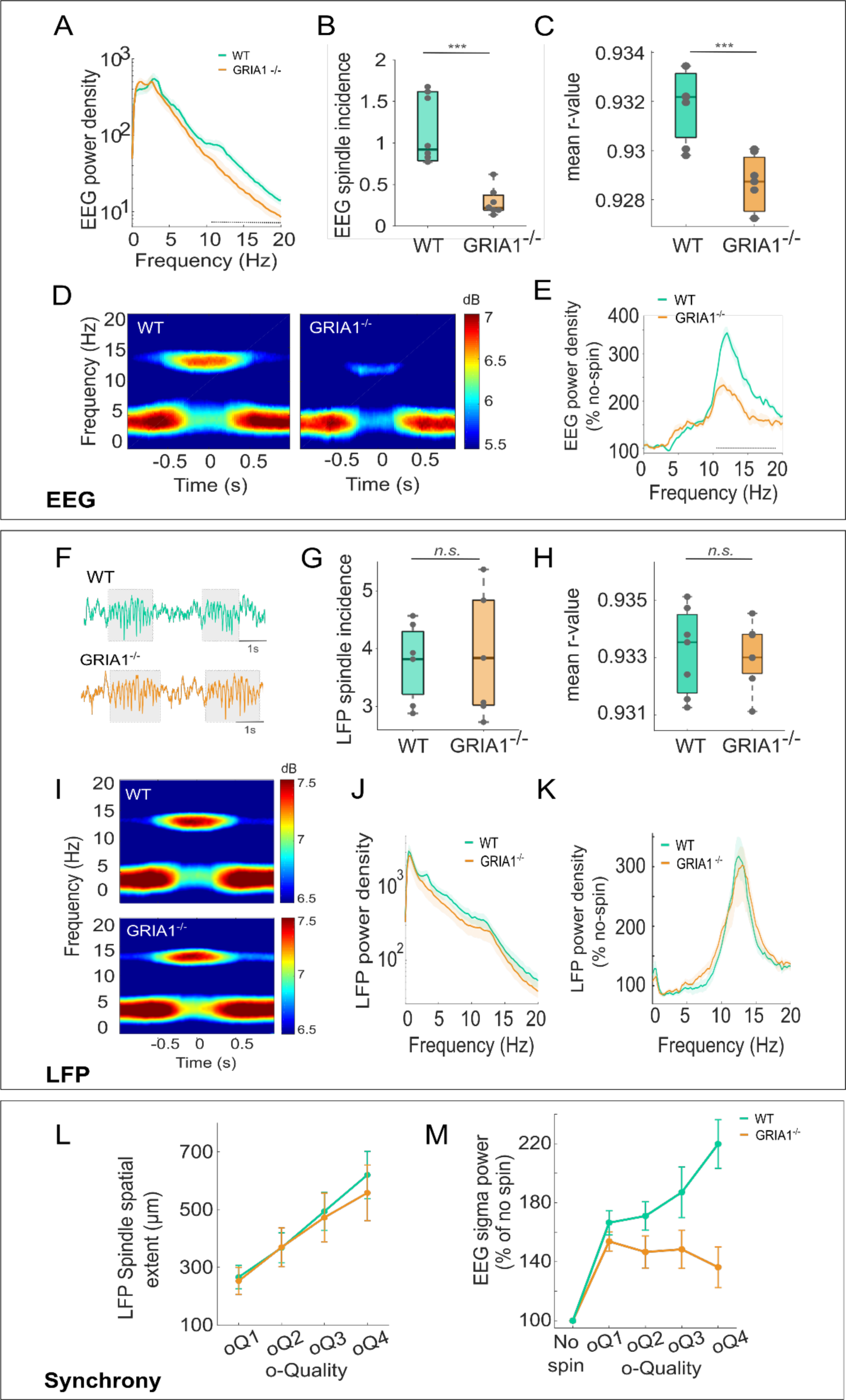
Glutamatergic neurotransmission is essential for large-scale but not local dynamics of spindles. **(A)** Mean EEG power density spectra during NREM sleep in the frontal derivation in GRIA1-/- mice and WT littermates. Shaded area= SEM. **(B)** Mean frontal EEG spindle incidence per minute of NREM sleep for GRIA1-/- mice and WT littermates. **(C)** Mean maximum r-value for EEG spindles detected in the frontal EEG in GRIA1-/- mice and WT littermates. **(D)** Spectrograms centered around the midpoint of individual spindle events in the frontal EEG channel recorded from WT (top) and GRIA1-/- (bottom) mice. EEG spectral power represents mean across mice, where GRIA1-/- (n=7) and WT mice (n=5). Spectrograms are colour-coded on a logarithmic scale (dB). **(E)** Mean EEG power density spectra in the frontal EEG channel during epochs with automatically detected spindle events. Mean (lines) and SEM (shaded area) are shown for each frequency bin expressed as a percentage of mean power density during NREM sleep epochs without detected spindles (‘% no spin’). Genotype*frequency: F_80,560_=6.47, *p*<0.001 (*GG*); effect of Genotype: F_1.3,4.1_=14.79, *p*<0.001 (*GG*). **(F)** Six second segments of LFP signal (500μV amplitude range) recorded from layer 4 of S1 from one WT and one GRIA1-/- mouse, showing illustrative examples of LFP spindle events in both genotypes. Detected spindles are highlighted with grey shading. **(G)** Mean spindle incidence per minute of NREM sleep for LFP signals recorded from layer 4 in GRIA1-/- mice and WT littermates. **(H)** Mean maximum- r value for LFP spindles detected in layer 4 in S1 from GRIA1-/- mice and WT littermates. **(I)** Spectrograms centered around the midpoint of individual spindle events in a channel located in layer 4 of S1 channel recorded from WT (top) and GRIA1-/- (bottom) mice. LFP spectral power represents mean across mice, where GRIA1-/- (n=7) and WT mice (n=5). Spectrograms are colour-coded on a logarithmic scale (dB). **(J)** Mean LFP power density spectra during NREM sleep in the LFP signal recorded from layer 4 of the S1 cortex in both GRIA1-/- and WT mice. The shaded areas represent the SEM. **(K)** Mean LFP power density in a channel located in layer 4 of S1 during epochs with automatically detected spindle events. Mean (lines) and SEM (shaded area) are shown for each frequency bin expressed as a percentage of mean power density during NREM sleep epochs without detected spindles (‘% no spin’). Genotype*frequency: F_80,960_=0.26, p=0.74; effect Genotype: F_1,11_=0.16, p=0.70. **(L)** Mean spatial extent of LFP spindles recorded from 16-channel micro-wire arrays from S1 in WT and GRIA1-/- mice, as a function of *o-Quality*. The spatial extent of a spindle is calculated from the number of LFP electrodes involved in each spindle event. **(M)** Mean EEG sigma power in the frontal derivation during epochs with spindles detected in the LFP (S1 – layer 4) as a function of the *o-Quality* of LFP spindles in both genotypes. Mean (lines) and SEM (error bars) are shown expressed as a percentage of mean power density during NREM sleep epochs without detected spindles (‘% no spin’). (*Note:* EEG: electroencephalogram. WT: wild-type. SEM: standard error of the mean. LFP: local field potential. S1: primary sensory cortex. GG= Greenhouse-Geisser. Figures show mean and SEM across mice. Dotted lines depict bins where values differed significantly between genotypes. For boxplots: black lines= mean across mice, boxes= SEM, whiskers= 95% confidence intervals, dots= individual values for each mouse. GRIA1-/- (n=7) and WT mice (n=5). *p<0.05, **p<0.01, ***p<0.001.).

Unexpectedly, visual inspection of LFP signals in S1 revealed the occurrence of well-defined NREM spindle events in S1 in all individual GRIA1^-/-^ mice (Fig.6F). These events were characterised by a similar incidence (t_12_=0.32, *p*=0.71) and *o-Quality* (t_12_=0.17, *p*=0.86) as in WTs (Fig.6G-H) and were associated with comparable levels of spectral power in the corresponding LFP signal (Fig.6I-K, Genotype x frequency: F_80,960_= 0.23, *p*=0.77; Effect of genotype: F_1,12_=1.35, *p*=0.27). Additionally, in both genotypes there was a positive relationship between the *o-Quality* and spatial extent of LFP spindles (Fig.6L, Genotype x *oQ*: F_1.15,13.86_=1.96, *p*=1.84. Effect of *o-Quality* on spindle spatial extent F_1.15,13.86_=91.5, *p*<0.0001.), suggesting that the local synchrony of spindles in S1 is intact in GRIA1^-/-^ mice. However, in the GRIA1^-/-^ mice the occurrence of S1 LFP spindles, even of high *o-Quality*, was only weakly associated with any changes in the frontal EEG (Fig.6M, Genotype x *oQ:* F_1.9,22.9_=13.89, *GG*, *p*<0.0001; KO F_1.19,7.18_=1.69 *GG*, *p*=0.24; WT F_1.96,11.76_=14.98 *GG*, *p*<0.001). One interpretation of this finding is that the deletion of the GluA1 AMPA receptor subunit results in a failure of S1 spindles to propagate to distant cortical areas. This finding suggests an important role of GluA1-mediated neurotransmission or synaptic plasticity in the regulation of large-scale network synchronisation of sleep spindles.

### The o-Quality of spindles is inversely related with the behavioural responsiveness to auditory stimulation during sleep

Mounting evidence suggests that spindles protect sleep from sensory disruption^30–38^, possibly by reducing the relay of sensory information in the thalamo-cortical network^124–126^. As our results indicate that the spindle *o-Quality* metric reflects synchrony within the thalamo-cortical network, we hypothesized that the *o-Quality* of spindles will inversely correlate with the responsiveness to auditory stimulation during sleep. One approach to measure the degree of sensory disconnection is to measure the motor response to sounds played during sleep^127, 128^. Here we quantify motor responses as the variance of the EMG signal recorded from the nuchal muscle. To this end, we developed a real-time event-triggered stimulation system, which allowed us to deliver online auditory stimuli during the presence or absence of spindle events detected in S1 (Fig.7A), the brain area where spindles are most prominent^50, 51^, and assess the effects of this stimulation on the EMG response as a readout of sensitivity to the auditory stimulus. Spindles were detected in real-time based on the sigma power calculated online from LFP signals recorded from layer 4 of S1 (Fig.S6A-B).

**Fig. 7.**
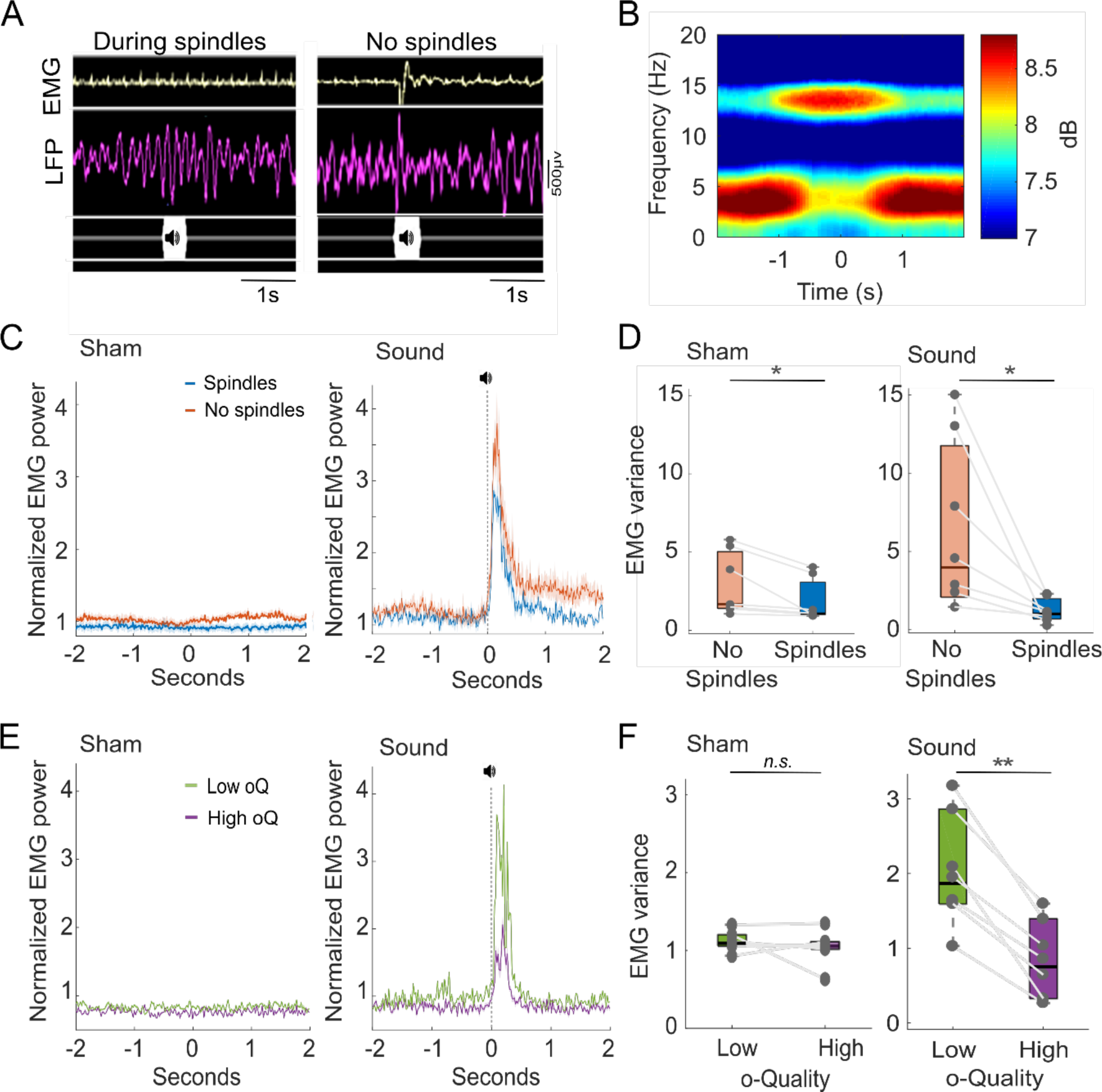
The o-Quality of spindles is inversely related with the behavioural responsiveness to auditory stimulation during sleep. **(A)** Examples of auditory stimulation during (left) and outside (right) spindle events, showing 3-s LFP and EMG segments for one mouse. **(B)** Spectrogram centred around the midpoint of individual spindle events detected in an LFP signal recoded from layer 4. LFP spectral power represents mean across mice (n=7). Spectrograms are colour-coded on a logarithmic scale (dB). **(C)** EMG response to sham stimulation (left) and sound stimulation (right) at time 0, showing stimulations delivered during spindles (n=1400) and stimulations in non-spindle NREM sleep (n=1700). **(D)** EMG variance during the 200 ms period of sham stimulation (left) and auditory stimulation (right) delivered outside or during spindle events. **(E)** EMG response to sham stimulation (left) and sound stimulation (right) at time 0, showing stimulations during spindles of high (oQ4) (n=170) and low (oQ1) (n=780) *o-Quality*. **(F)** EMG variance during the 200ms period of auditory stimulation (left) or sham stimulation (right) delivered during spindle events of with high (oQ4) and low (oQ1) *o-Quality.* (*Note:* In C-F the EMG power (µV^2^) is normalized to the EMG power (µV^2^) during NREM epochs with no stimulation. LFP: local field potential. EMG: Electromyography. Lines= average across mice, shaded area=SEM. For boxplots: black lines= mean across mice, boxes= SEM, whiskers= 95% confidence intervals, dots= individual values for each mouse. *p<0.05, **p<0.01, ***p<0.001).

As expected, spectrograms centred at the time stamp of a real-time detection of individual spindles, showed a prominent increase in LFP power within the spindle frequency range in S1 (Fig.7B). The comparative sensitivity (comparative true-positive rate) between detections made with the real-time detector and offline detections with the lowest threshold of the AR-model (r*_b_*=0.92), reached 86.2% ± 2.11. Auditory stimulation was presented from ZT3.5 to ZT9.5 and consisted of brief (100ms) pure tones (10KHz) played at either 70dB (‘sound condition’) or 0dB (‘sham condition’) (Fig.S6C-D). The sound and sham conditions were presented on two different days and the order of presentation was counterbalanced across mice.

We first confirmed that the sound stimulation did not affect the properties of spindle events or the distribution of vigilance states across the 12h light-period. Specifically, we found that the percent of time mice spent in NREM (F_11,66_=0.74, *p*=0.69), REM (F_11,66_=1.1, *p*=0.41,) and wake (F_11,66_=0.87, *p*=0.60) states, or the number of brief awakenings (F_11,66_=1.43, *p*=0.23) did not differ between the sound and the sham conditions (Fig.S7A-C). Similarly, we found that the incidence (t_6_=0.43, *p*=0.68), duration (t_6_=0.16, *p*=0.88) and *o-Quality* (t_6_=0.27, *p*=0.79) of spindles did not differ during sound relative to sham stimulations (Fig.S8A-D). Similarly, spindle frequency changed by only 0.1Hz (t_6_=4.38, *p*<0.01) between conditions (Fig.S8C). LFP power at the sigma frequency range also did not differ with sound relative to the sham stimulation (z-test=0.06, *p*=0.99) (Figure.S8E).

Next, we calculated the EMG response to auditory stimulation (i.e. sound vs. sham) across spindle conditions (i.e. present vs. absent). Considering epochs with sham stimuli (0dB) only, we found that the EMG variance was significantly lower when spindles were present as compared to trials when spindles did not occur (mean difference = 1.09; F_1,6_=7.69, *p*<0.05) (Fig.7C-D). This indicates that muscle activity is generally lower during spindle events. Notably, this mean difference in EMG variance was even greater during the ‘sound stimulation’, when auditory stimulation (70dB) was delivered at the time of spindles or outside spindle events (mean difference = 5.86, F_1,6_=7.12, *p*<0.05), leading to a significant spindle condition (present vs. absent) x stimulation (sound vs. sham) interaction (F_1,2_=4.7, *p*<0.05) (Fig.7D). These results suggest that overall, the EMG response is lower at the time of spindle occurrence and the presence of spindles in S1 is related to an attenuated EMG response to auditory stimulation.

Next, we investigated whether the EMG response to auditory stimulation varied in relation to the *o-Quality* of S1 spindles detected offline with the AR-model. We found that the variance of the EMG signal was significantly higher when sounds were played during spindles with low *o-Quality* relative to spindles with high *o-Quality* (F_1,6_=36.11, *p*<0.01, Fig.7E-F). Importantly, this difference was not present during the sham stimulation condition (F_1,6_=0.2, *p*=0.89, Fig.7E-F). These results suggest that the *o-Quality* of spindles is inversely related with the behavioural responsiveness to auditory stimulation during sleep. This is in line with the hypothesis that spindles protect sleep from sensory disruption, but importantly, these findings highlight that not only the presence but also the *o-Quality* of spindles provides relevant insights into their functional role.

## Discussion

The primary aim of this study was to investigate the origin and the functional significance of the variability and heterogeneity among sleep spindles, in terms of their characteristics and spatio-temporal dynamics. As well known, a wide range of neurophysiological parameters are best described with a lognormal distribution^129^, and we now demonstrate that this includes the fundamental defining properties of sleep spindles, such as their damping and amplitude. The view that sleep spindles are discrete, all-or-none events has dominated the field for decades and has been instrumental in understanding their neurophysiological mechanisms and functions^54–56^. However, the time is ripe to acknowledge that that there is no mathematical or physiological justification for defining a single meaningful threshold for separating spindle events from background sigma activity. Rather than a methodological inconvenience, we thought that incorporating this property of spindles into their definition will not only advance our understanding of their underlying neurophysiological mechanisms but will allow to bridge the gap between the questions “how come spindles occur?” and “what they are for?”.

This requires the development of a novel metric, and to that end, we propose here the concept of oscillatory-Quality (*o-Quality*), to measure and parameterize the strength of oscillatory events occurring at the spindle frequency range (10-15Hz). The *o-Quality* metric is derived by fitting an auto-regressive model to short segments of electrophysiological signals and using it to identify and calculate the damping of spindle oscillations. We found that the *o-Quality*: (1) captures a wide range of spindle properties related to their spatio-temporal dynamics; (2) directly reflects the degree of network synchronisation; (3) correlates with the probability of spindle-slow wave coupling, and (4) is inversely related to the behavioural responsiveness to auditory stimulation during sleep. These findings, together with the observations that *o-Quality* of spindles is sensitive to manipulations targeting glutamatergic neurotransmission and preceding sleep-wake history, points to the global regulation of synaptic strength as one of its possible neurophysiological substrates.

### The o-Quality of sleep spindles is an emergent property of their spatiotemporal dynamics

The present study supports previous findings, which suggest that many characteristics of spindles are strongly influenced by the topography of their occurrence^42–50, 69^. Furthermore, we find that *o-Quality* of sleep spindles varies substantially between microscopic and mesoscopic regions, and this variability shows distinct topographic gradients. At the EEG level, the spindle *o-Quality* is higher in anterior cortical areas, while intra-cortically the *o-Quality* of spindles was higher in anterior regions of S1 and comparatively lower in M1. However, our results also demonstrate that within a specific cortical region (i.e. S1), one fundamental laminar profile describes all spindles regardless of their *o-Quality*, suggesting that spindles with different *o-Quality* have similar generating networks. Additionally, we demonstrate that the laminar profile of spindles shows regional variations, which is consistent with previous findings showing that the density of thalamo-cortical projections to different layers varies across cortical areas^130^. In line with previous studies^66, 67^, we found that the incidence and *o-Quality* of spindles is higher in layers 3 and 4 of S1. While in M1, where thalamo-cortical projections form the majority of synapses in layer 5^130^, spindle incidence and *o-Quality* is highest in the superficial subdivision of layer 5 and layer 4.

### The o-Quality of sleep spindles reflects network synchronisation

Overall, our results support the view that spindles are primarily local phenomena^43, 45, 50, 131^, but it is also common that spindles can occur across large cortical regions^82, 132–135^. Moreover, our results indicate that *o-Quality* of sleep spindles reflects the levels of synchronisation within and across cortical networks. Specifically, spindles with low *o-Quality* are typically observable within a few nearby recording sites only, and are transient, while high *o-Quality* spindles persist longer and encompass larger cortical regions.

Importantly, our results show that the vast majority of local LFP spindles remain undetected at the global EEG level. This suggests that studies of spindle dynamics based on “global” EEG recordings should be viewed with caution as these include only a small proportion of high *o-Quality* and synchronous events, omitting the majority of local spindles. Of course, at present, it is not feasible to obtain intracranial recordings in humans outside of clinical contexts. Nonetheless, these results indicate that high *o-Quality* sleep spindles reflect, in general, a more synchronised state of cortical networks.

To further address the relationship between spindle *o-Quality* and network states, we tested the hypothesis that oscillatory strength of spindles will correlate with their coupling with other sleep oscillations, such as slow-waves. Consistent with this hypothesis, we found that high *o-Quality* spindles are more likely to be preceded by high-amplitude LFP slow waves than low *o-Quality* spindles. These results are in line with findings suggesting that slow waves are involved in entraining spindle events^66, 82–87, 136^. The concurrent increase in SWA and spindle *o-Quality* after sleep deprivation also support the idea that sleep need is associated with a more efficient recruitment of large neuronal populations in network oscillations^91^. This notion was supported by the observation that the *o-Quality* is a reliable measure of how strongly neuronal spiking is modulated during spindle oscillations. Taken together, these results suggest that the *o-Quality* of spindles reflects synchrony within cortical networks, which is sensitive to the levels of homeostatic sleep need.

Previous studies have shown an inverse correlation between sigma and slow-wave activity across the light period (12-h)^47, 98^ and during the first hours of recovery sleep after sleep deprivation^47, 93, 96, 97^. Our results show, however, that the association between SWA and spindles varies based on the spindle *o-Quality*. Specifically, while the occurrence of low *o-Quality* spindles (which show higher overall incidence) shows the typical negative correlation with SWA across the 12h light period, high *o-Quality* spindles show a positive correlation with SWA. These results raise an interestingly possibility that *o-Quality* of sleep spindles may be informative about the state of cortical networks in general, beyond being merely a metric specific to sleep spindles only.

### The GluA1 subunit of the AMPA receptor is essential for large-scale but not local dynamics of spindles

To address the underlying neurophysiological mechanisms linking the network states with oscillatory dynamics of sleep spindles, we detected spindle events in transgenic mice deficient of the GluA1 AMPA receptor subunit^51^. These mice are an important model for investigating the role of synaptic plasticity in behaviour and sleep regulation^137^. Surprisingly, we observed that despite a profound reduction in the incidence and *o-Quality* of EEG spindles in the frontal cortex, LFP spindles in S1 were preserved in GRIA1^-/-^ mice. Furthermore, despite these S1 spindles showing comparable *o-Quality* in WT and GRIA1^-/-^mice, they completely failed to express in distant cortical areas in the animals lacking the GluA1 subunit.

While the exact mechanisms underlying these striking effects remain to be determined, our findings shed new light on the origin and dynamics of sleep spindles. First, they suggest an important, and hitherto under-investigated link between glutamatergic neurotransmission and the network mechanisms implicated in the generation and propagation of spindles. AMPA and NMDA receptors are known to play an important role in the generation of thalamo-cortical oscillations^138, 139^, but the nuanced role of GluA1 subunit specifically has not been previously recognised. Crucially, we find that the deletion of this subunit does not affect the capacity to generate spindles or the persistence of spindle-activity within local cortical networks. Instead, it primarily affects the large-scale network synchronization of spindle-activity, as reflected in spindle events remaining localised and virtually undetectable merely a few millimetres away from the site where they are prominent.

In line with this hypothesis, electron microscopy evidence suggests that although thalamo-cortical and corticothalamic synapses in the reticular nucleus of the thalamus express high levels of AMPA receptors, these contain mainly GluA4 and some GluA2/3 subunits. The GluA1 subunit, however, is barely detectable in this brain region^140^. In contrast, GluA1-rich AMPA receptors are expressed in high levels in synapses between thalamo-cortical projecting cells and fast-spiking interneurons in the cortex. It has been suggested that among other mechanisms, these GluA1-rich AMPA receptors could provide rapid activation kinetics capable of recruiting feedforward inhibitory circuits that could propagate spindles across cortical circuits^141^. In fact, there is evidence suggesting that spindle network synchrony is regulated by intracortical connectivity and cortico-thalamic feedback control^19, 66, 135, 142^.

These results also have potential clinical implications given the link between GluA1 and neuropsychiatric disorders like schizophrenia. Several studies have reported that EEG spindles are reduced in patients with schizophrenia^113–123^, and further suggest this may reflect deficits in the thalamic reticular nucleus in this disease^114, 116, 122, 143^. Our results suggest the intriguing possibility that large-scale synchronization deficits, resulting from the disruption of glutamatergic pathways, could alter the expression of spindles at the global EEG level in schizophrenia, even when (at least some) spindle initiation mechanisms are preserved. It is also possible that these spindle disruptions may contribute to the fragmented sleep^144^ or memory deficits^117, 145^ reported in patients with schizophrenia. Electrophysiological recordings across cortical layers combined with recordings or manipulations of the reticular nucleus of the thalamus, would be relevant to further understand the association between GluA1, cortico-thalamic feedback control, and spindle network synchrony.

Finally, given the important role of GluA1 subunit in the mechanisms of synaptic plasticity^103, 146^, we cannot exclude the possibility that the emergence and propagation of spindle-activity during sleep depends on how strong or efficacious the synapses are across the cortex or thalamo-cortical networks. The functional role of sleep spindles in offline information processing, memory replay or synaptic renormalisation has received considerable attention in the last decades^24, 141, 147–150^. Our data now suggest an intriguing possibility that spindles are, in turn, regulated by the levels of synaptic strength or the capacity to modify synaptic efficacy, possibly in a sleep-dependent manner, which opens an entirely new avenue into understanding their functional role.

### The o-Quality of spindles is inversely related to the behavioural responsiveness to auditory stimulation during sleep

Evidence suggests that sleep spindles may support the maintenance of sleep by disrupting the transfer of sensory information to the cortex^30–38^. Nevertheless, the neurophysiological mechanisms underlying this effect remain unclear, and there is also contradictory evidence for this notion^151^. In our study, we found that motor responses (measured as EMG variance) to auditory stimulation are significantly reduced when stimuli are delivered during spindles compared to NREM sleep in the absence of spindles. Importantly, our findings further suggest that not only the presence, but also the *o-Quality* of spindles matters, as the magnitude of motor responses to auditory stimulation presented during spindles showed an inverse relationship with the spindle *o-Quality*. Spindles with high *o-Quality* are related to a reduced responsiveness to auditory stimulation during sleep, which suggests increased sleep protection.

The potential role of spindles in protecting sleep from environmental disruption has been attributed to the thalamic origin of these oscillations. The thalamus relays sensory information to the cortex^125^ and is an important control centre that shapes sensation and action^152^. These operations require precise inhibitory control, which is largely driven by innervation from structures like the reticular nucleus of the thalamus^16, 124, 153, 154^. It has been shown that burst firing generated during spindles can quench these sensory inputs^124, 126^. Specifically, this burst firing reduces the action potential output that thalamo-cortical neurons generate relative to their excitatory input. This has been proposed as one of the mechanisms through which burst firing in thalamo-cortical networks, which gives rise to oscillations like spindles, could reduce the transfer of sensory information to the cortex during sleep^147^.

In line with these hypotheses, human studies have shown that sensory stimuli fail to generate evoked responses in the cortex and need to have increased intensity to wake subjects when stimulation occurs in phase with spindle events detected in the thalamus or cortex^30, 34, 35, 155^. Additionally, the density of EEG spindles during spontaneous sleep positively correlates with the tolerance shown by subjects to environmental noise during sleep^30^. Combined EEG and fMRI studies in humans, have also shown that pure tones elicit brain responses in the thalamus and primary auditory cortex, which is similar during NREM sleep and wake. These brain responses in the thalamus and the primary auditory cortex are reduced or absent when the sounds are paired with spindles or the down-states of the slow oscillation^32, 156^. Additionally, mice over-expressing Ca^2+^-dependent small-conductance-type 2 potassium (SK2) channels (which have been found to support spindle generation), show enhanced thalamic spindle activity together with decreased responsiveness to noise exposure during sleep^36^. Our results are in line with these previous findings and further indicate that the spindle *o-Quality* metric reflects synchrony within the thalamo-cortical network and the *o-Quality* of spindles affects the responsiveness to auditory stimulation during sleep.

## Conclusions

In this study we develop and validate a novel quantitative metric to characterise sleep spindles in terms of their oscillatory strength. We further provide abundant evidence that the *o-Quality* of sleep spindles reflects many fundamental properties of spindle activity – from their topographical and laminar distribution, to their involvement in sensory processing during sleep, coupling with other network oscillations and sleep homeostasis. The most important, and provocative, implication of our study is the demonstration of how developing a new approach to describe a neurophysiological phenomenon can pave the way for understanding its functional meaning. Shifting attention from reporting how a specific experimental intervention affects “quantity” of sleep spindles to their *oscillatory-Quality*, in our view, represents a major step forward, which, without doubt, will bring us closer to providing a better mechanistic understanding of brain oscillations in health and disease.

## Methods

### Animals

Experiments were performed in adult male C57BL/6 mice (n=34) and adult male GRIA1^-/-^ (n=7) and littermate WT (n=7) mice (mean age 16.9±0.5 weeks and mean weight 32.5g ± 2.1g [mean ± SEM] at the time of experiments). All mice were bred at the Biomedical Sciences Building (University of Oxford, UK). GRIA1^-/-^ mice were generated as previously described^99^ and maintained on a C57BL/6J x CBA/J background. Heterozygote parents were bred, resulting in ∼25% GRIA1^-/-^ mice that lacked both copies of the GluA1 allele, ∼25% WT mice that had both copies of this allele, and ∼50% heterozygote mice that were not used here. At the end of all experiments, the genotype of mice was confirmed by genotyping. This was carried out by TransnetYX, USA, using ear notch samples and PCR-mediated amplification methods. During the experiments, mice were individually housed in plexi-glass cages (20.3 x 32 x 35cm) under a 12:12 light-dark cycle (lights on at 9 am). Cages were housed in sound-attenuated, electro-magnetic shielded, ventilated Faraday chambers (A Lafayette Instrument Company, USA). Food and water were available *ad libitum*. Room temperature and relative humidity were maintained at 22 ± 1°C and 60 ± 10%, respectively. Experimental procedures were performed in accordance with the Animal (Scientific Procedures) Act 1986 under a UK Home Office Project Licence (P828B64BC) and were in accordance with institutional guidelines.

### Surgical Procedure and electrode configuration

Surgical procedures were performed under isoflurane anaesthesia. All mice (n=48 in total) were implanted with epidural screws to record EEG signals, intra-cortical probes to record local field potentials (LFP) and multi-unit activity (MUA), and tungsten wires in the nuchal muscle to record electromyography (EMG). EEG/EMG mounts were composed of stainless-steel screws (shaft diameter 0.86mm, InterFocus Ltd, UK) and two single stranded stainless-steel wires, attached to an 8-pin mount connector (8415-SM, Pinnacle Technology Inc, USA) as described previously^157–161^. EEG screws were implanted epidurally over frontal (+2mm anteroposterior (AP), +2mm mediolateral (ML), relative to bregma), parietal (−0.5 to −1.5 mm AP, 2mm ML) and/or occipital (−4mm AP, 2.5 mm ML) cortical regions (Fig.S9). A reference screw was implanted over the cerebellum and an anchor screw was implanted contralaterally to the EEG screws to provide stability to the implant. Finally, the EMG was recorded from the two stainless steel wires inserted on both sides of the nuchal muscle. All the screws and wires were attached to the skull using dental cement.

LFPs and MUA were recorded across or within cortical layers using two different types of electrode arrays (Fig.S9). To record signals across layers of the cortex, mice were implanted with 16-channel laminar probes (NeuroNexus, A1×16-3mm-100-703, 100μm spacing) either in the anterior area of primary somatosensory cortex (S1, n=7 mice, +0.3mm AP and −3.25mm ML), a more posterior area of S1 (n=7 mice, −0.7mm AP and −3.25mm ML) or the primary motor cortex (M1, n=7, 1.1mm AP and −1.75mm ML). In a subset of animals (n=7 C57/BL6; n=7 GRIA1^-/-^; n=7 WT littermates), a polyimide-insulated tungsten microwire array (Tucker-Davis Technologies Inc, USA) was implanted into deep layers of S1 (layers 4-5), with recording tips positioned approximately equidistant to the cortical surface in the anterior-posterior direction (where well-defined spindles have been previously reported in mice) ^50, 162^. Microwire arrays consisted of 16-channels with properties as follows: two rows of eight wires, wire diameter 33μm, electrode spacing 250μm, row separation L-R: 375μm and tip angle 45 degrees. Arrays were customised so the left row was 250μm longer. For microwire arrays recording, a craniotomy of approximately 1×2mm was made and the midpoint of the craniotomy was located relative to bregma: −1mm AP and −3.25mm ML.

### Histological verification of recording site

To confirm the location of the electrodes (Figure S1), all laminar probes and micro-wire array wires were coated with DiI fluorescent dye (DiIC_18_(3), Invitrogen), prior to their implantation. At the end of the experiment, mice were deeply anaesthetised, electrolytic micro-lesions (10μA, 20s) were performed at specific sites to be used as landmarks to verify the recording locations (NanoZ, Neuralynx), and mice were transcardially perfused (0.9% saline, 4% paraformaldehyde) as described previously^157^. Brains were sliced to obtain 50μm coronal slices, using a Vibratome (Leica VT1000 S, Germany). The brain slices were stained with DAPI, mounted on slides, and imaged with a fluorescence microscope (Olympus Bx51, Japan), using 1.6, 2.5, and 5x magnifications. The electrode locations were mapped using the Dil stain, microlesion traces and the coordinates of the recording sites were identified using a mouse brain atlas ^163^. The depth of the implants was assessed measuring the distance between the cortical surface and the electrical current induced tissue microlesions^157^. ImageJ (v1.52a) was used to merge fluorescence images and add scale bars^164^. All figures were created using Inkscape (v1.0.2, Inkscape Project 2020; https://inkscape.org).

### Signal processing and analysis

Electrophysiological recordings were acquired with a RZ2 High Performance Processor and Synapse software (Tucker-Davis Technologies Inc., Alachua, FL, USA). EEG, EMG and LFP signals were continuously recorded, concomitantly with extracellular neuronal spike data from the same electrodes used for LFP monitoring (PZ5 NeuroDigitizer pre-amplifier, TDT, USA).

EEG, LFP and EMG signals were filtered between 0.1–100Hz and stored at a sampling rate of 305Hz. The signals were resampled offline at a sampling rate of 256 Hz using custom-made Matlab (The MathWorks Inc, Natick, Massachusetts, USA) scripts. For subsequent analyses, EEG and LFP power spectra were computed by a Fast Fourier Transform of 4-s epochs (Hanning window), with a 0.25Hz resolution Matlab (The MathWorks Inc, Natick, Massachusetts, USA).

Extracellular neuronal activity was continuously recorded at a sampling rate of 25kHz and filtered between 300Hz - 5kHz. For spike acquisition, amplitude thresholds were manually set on Synapse on each recording channel^157, 165^. This threshold was set at least 2 standard deviations above noise level. When the recorded voltage crossed this predefined set threshold, 46 samples around the event (0.48-ms before, 1.36-ms after the threshold crossing) were extracted (Fig.S10). Spike waveforms were processed using custom-made Matlab scripts. Events with artefactual waveforms were excluded from further analysis^157, 161, 165^.

### Scoring and analysis of vigilance states

Sleep scoring was performed offline and manually (Fig.S10). Resampled EEG, LFP and EMG signals (256Hz) were transformed into European Data Format (EDF) using open source EEGLAB (Swartz Center for Computational Neuroscience, La Jolla, California, USA). These signals were visualized in 4-s epochs using the software SleepSign (Kissei Comtec Co, Nagano, Japan). Wake was defined as low voltage, high frequency EEG activity accompanied by a high level of EMG activity lasting more than 4 epochs. NREM sleep was defined as signal with high voltage and slow frequency, predominantly characterised by the occurrence of slow-waves (0.5-4Hz) and sleep spindles (10-15Hz). REM sleep was defined as low voltage, high frequency oscillations, with predominance of theta (6-9 Hz) activity in occipital derivations, which was distinguished from waking by the reduced level of EMG activity. Brief awakenings (microarousals) were defined as transient periods of low voltage, high frequency oscillations in the EEG and LFP signals accompanied by elevated EMG tone, lasting ≥4s and ≤16s (Fig.S10). Epochs containing artefacts, resulting from eating, drinking or gross movements, were identified and removed from the analyses. Overall, 11.2%±2.9% of wake, 0.6%±0.8% of NREM and 0.8%±0.7% of REM epochs contained artefactual EEG and/or LFP signals across all animals.

### Spindle detection

Oscillatory events were detected in all EEG and LFP signals by applying a previously published algorithm^10, 12, 14^, based on autoregressive (AR) modelling of the EEG. This algorithm models electrophysiological brain signals as a superposition of stochastically driven harmonic oscillators (*f* > 0Hz) and relaxators (*f* = 0Hz) with damping and frequency varying in time. For this analysis, filtered (0.1–100Hz) EEG and LFP times-series *x(t)* were resampled at 128Hz and overlapping 1-s segments, shifted by 1 sampling interval, were modelled with an autoregressive model (AR-model) of order *p=*8. As such, the AR(8)-model uses the weighted sum of the preceding *p* samples to predict the value of the *n^th^* sample of the time series *x*(*t*_*n*_):

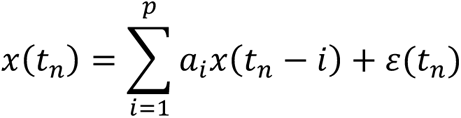

where *a*_i_ denotes the AR coefficients and *ε*(*t*_*n*_) the residuals. The model was estimated using the Burg algorithm^166^. These *a*_i_ coefficients are related to the frequency *f*_*k*_ = ∅_*k*_ (2*π*Δ) and damping coefficient 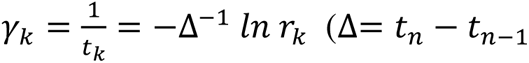 denotes the sampling interval) using:

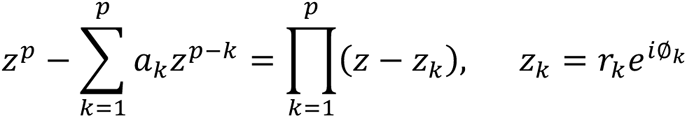

Note that *r*_*k*_ is exponentially related to the damping coefficient *γ*_*k*_, such that a decrease in *γ*_*k*_ is reflected as an increase in *r*_*k*_. Additionally, the order of the AR-model (*p*) determines the total number of poles *z*_*k*_ (by *p=2m + n,* with *m* oscillators and *n* relaxators), such that our AR(8)-model could generate up to four different oscillatory poles *z*_*k*_. In this way, when the signal is dominated by a rhythmic and stable oscillation with frequency *f*_*k*_, like a spindle (10-15Hz), this activity will be reflected by a reduction in damping coefficient *γ*_*k*_and an increase in *r*_*k*_ of the corresponding pole *z*_*k*_ with the frequency *f*_*k*_ (Fig.2A-D).

Two types of thresholds were used to detect oscillatory events – named here the upper and the lower one. As described previously^10, 14^, oscillatory events were detected when the damping coefficient *γ*_*k*_ at frequency *f*_*k*_decreased and, therefore, *r*_*k*_surpassed a predefined upper threshold *r*_*b*_. To this end, we have chosen the threshold of 0.92, initially based on the visual inspection of multiple representative recordings in each animal, which revealed that spindle oscillations (10-15Hz activity) were not easily discernible from background activity when *r*_*b*_was lower than 0.92. To further validate the choice of this threshold, surrogate EEG and LFP signals (see *surrogate signal generation* section below) were created from the original signals recorded during NREM sleep for every animal and derivation. Next, we calculated the distribution of r-values for oscillators at frequencies between 10-15Hz for both the original signals and their respective surrogates. We then calculated the ratio between the r-values of real and surrogate signals (Fig.S11). In all cases, the percentage difference between real and surrogate signals was at least 92%, corresponding to a r*_b_* level of 0.92. We acknowledge that this approach is somewhat arbitrary, and the choice of the upper threshold value should be considered operational, which, however, is in line with the key conclusion of our study that spindles are not all-or-none events.

For each event, the start time (*t_1_*) was considered the time point when *r*_*k*_ exceeded *r*_*b*_=0.92, while end time (*t_2_*) was considered the time point when *r*_*k*_ fell below *r*_*b*_=0.92. Subsequently, we grouped detected oscillatory events into four groups, named oscillatory quality (*oQ*) 1 to 4 (*oQ*1 to *oQ*4), corresponding to r-values between 0.92-0.95 (i.e. high-to-low damping), with an additional group including all events above r=0.95 (Fig.2E). This approach allowed us either to quantify the incidence of oscillatory events within a specific *oQ* group, or investigate the effects of experimental manipulations, brain regions or cortical layers on the average *oQ*-value across all detected events within the relevant time interval.

The lower threshold *r*_*a*_ was used to address transient fluctuations of *r* and to merge or split overlapping or immediately following events within a specific frequency. Specifically, consecutive oscillatory detections were considered a single continuous event if *r*_*k*_ stayed above the lower threshold *r*_*a*_, or were considered separate events if *r*_*k*_ fell below *r*_*a*_. Similar to previous applications^12, 14^, we set the lower threshold to *r*_*a*_=0.90.

The oscillatory events detected by this algorithm in the human sleep EEG correspond to the classically defined EEG frequency bands: e.g. delta (1.5 −4.5Hz), alpha (8-11.5Hz), and sigma (11-15Hz)^11, 14, 167^ (Fig.S2A). Detected oscillatory events that showed a mean frequency *f*_*k*_ between 10-15Hz were defined as putative ‘spindle events’. Figure 2 illustrates the principle of the algorithm and demonstrates the detection of several spindle events in an 8-s segment of an LFP signal. Importantly, this spindle detection approach does not require signal filtering in any specific frequency band and does not assume any specific oscillatory waveform, which is relevant as filtering can considerably distort electrophysiological signals^57, 58^.

### Slow wave detection

Slow waves were detected in the EEG and LFP signals following the method presented in Viazovskiy, et al.^81^ Specifically, slow waves were detected in the signals after band pass filtering between 0.5-4Hz, using a phase-neutral (forward-backward) Chebyshev Type II filter^168^, with stopband edge frequencies at 0.3-8Hz. The parameters of the filter were optimized visually to obtain the maximal resolution of the wave shape, as well as to minimise intrusions of fast frequencies (i.e. spindles). Slow waves were detected as positive deflections in the signal, between two consecutive negative deflections (separated by at least 0.1-s)^72, 73, 169^. For our analyses, we selected slow waves with peak amplitude greater than the median amplitude detected across all slow waves because high-amplitude slow waves accurately reflect homeostatic sleep pressure^81^ and correspond to well defined neuronal OFF periods^160, 170^.

### Surrogate signal generation

To test the hypothesis that the detected spindle events and observed spindle dynamics were not obtained merely by chance, but rather represent true physiological phenomena, we created surrogate data (artificial signals) (Fig.S12A). These surrogate signals were created based on an improved version of the Iterative Amplitude Adjusted Fourier Transform (IAAFT) algorithm developed by Schreiber et al.^171^ Specifically, for each mouse and derivation we calculated a windowed Fourier transform for random 10-min segments of EEG and LFP signal recorded during NREM sleep. The resulting Fourier phases were randomized and the inverse of the Fourier transform was calculated. Then the amplitudes of the resulting time series were adapted to match the original amplitude distribution. This was done by rank ordering the resulting time series and replacing the data points with the data point of the original time series with the same rank. This procedure was performed in an iterative manner in order for the surrogate signals to achieve a closer match to the amplitude distribution and the power spectra of the original signals. The comparison of the sigma peak distribution and the infra-slow spectral dynamics of both real and surrogate signals show that indeed the original amplitude distribution of the time series was preserved (Fig.S12B) while the endogenous dynamics of spindle activity were not reflected in the surrogates (Fig.S12C), i.e. the surrogates did not show the previously reported^34, 172^ coordinated 0.02Hz oscillation of the sleep spindle band.

IAAFT surrogates test the null hypothesis that the data represent a stationary linear Gaussian process observed with a monotone, but potentially nonlinear, measurement function. In our case, we used the nineteen LFP (n=19) and EEG (n=19) surrogate signals to assess whether the spindle events detected in the real signal represented true characteristics of an underlying physiological system, or whether they could simply be described by a stationary linear stochastic process (i.e. were obtained by chance). The selected number of surrogates per signal (n=19) corresponds to a 5% significance level for a one-sided test. In other words, if the observable is larger than the value for all surrogates, the null hypothesis can be rejected (i.e. the detected events are not random).

### Current source density and multiunit activity analysis

The analysis on the laminar profile of spindles was done with custom-written Matlab and Python scripts, as well as IBM SPSS Statistics 27. Spindles detected simultaneously in different channels were considered to be co-occurrent (and thus representing the same, unique spindle event) if their centres, as defined from the damping analysis, occurred within 500ms of each other. The maximum *o-Quality* metric across channels was then assigned to each unique spindle event. In order to compare results across mice, every electrode was assigned a cortical layer (2/3, 4, 5 or 6) based on histology. Layer 1 electrodes were omitted from the analysis due to being absent in some mice. The effect of *o-Quality* on the number of simultaneous detections across layers was determined using a one-way ANOVA for every mouse separately.

Waveform averages as in Fig. 4D were calculated by averaging the unfiltered LFP and the current source density (CSD) across spindle events, time-locked to the trough of the maximum-envelope cycle in a layer 4-centered channel. The maximum-envelope LFP cycle peaks were aligned across spindle events using interpolation before averaging. For further analysis, the raw LFP signal was bandpass-filtered across the complete recording to a range of 10-15Hz using a 4th-order, zero-phase shift Butterworth filter. Spindle events were extracted from the filtered LFP signal in 5000ms epochs aligned around the spindle centre in the maximum *o-Quality* channel. The current source density (CSD) analysis was computed on the bandpass-filtered epoch and smoothed across channels using the cubic interpolation method ‘interp1d’ from the SciPy Python package. The amplitudes of the LFP and CSD signals in each channel were estimated using their respective 1-s root mean square (RMS) from the spindle centre. These values were subsequently averaged across all channels within a layer. Using the maximum signal envelope from the Hilbert transform in each channel yielded essentially identical results as using the RMS value (not shown). We used a repeated-measure ANOVA with layers as within-subject factors and *o-Quality* and between-subject factors to evaluate the effect of layer and *o-Quality* on the laminar profile of spindles, similar to Ujma et al.^68^, and performed this for every mouse. Finally, we performed a principal component analysis (PCA) on the laminar LFP and CSD amplitude profiles of all unique spindles in each mouse separately (Fig.S3.C).

The instantaneous spindle phase was obtained from the Hilbert transform of the filtered LFP in a layer 4-centered channel. The phase-amplitude coupling of the multi-unit activity (MUA) was calculated by extracting the spike times within 1-s of the spindle centre, from one electrode in each layer’s centre and assigning to each spike its corresponding spindle phase value (Fig. 4E). Circular statistics (mean angle, resultant vector length and Rayleigh’s test of uniformity) were performed using the Python ‘PyCircStat’ toolbox. The effect of *o-Quality* on the laminar profile of the spiking mean angle and resultant vector length was assessed with a two-way repeated measure ANOVA (with layers as within-subject factors and *o-Quality* and between-subject factors), after averaging the mean angles and vector lengths across spindles for each quality. As the mean firing angles were within a small range (<0.5 radians) and thus non-periodic, no circular statistics were employed for the ANOVAs on the pooled averages.

### Sleep deprivation

To investigate the effect of preceding sleep/wake history and sleep pressure on the characteristics of sleep spindles, slow waves and spectral parameters, total sleep deprivation was performed. This was done during the circadian phase when mice are typically asleep and therefore the homeostatic response to sleep loss can be reliably measured^79, 173^. To achieve sleep deprivation in an ecologically relevant manner, at light onset the nesting material was removed from the home cages, and mice were presented with novel objects to induce spontaneous exploratory behaviour. This intervention was performed the day following a 24-h undisturbed (baseline) recording. At the end of 6-h of sleep deprivation, all objects were removed, and the nesting material was returned to the cages. The procedure was successful, as mice spent only a minimal percent of the time asleep during sleep deprivation (1.19±0.42% of 6-h; n=14).

### Auditory stimulation based on real-time spindle detection

#### Real-time spindle detection

To date only a few studies have developed real-time spindle detection algorithms^174–176^, and fewer studies have performed real time acoustic stimulation triggered by spindle detections^174^. Here we developed and applied a real-time spindle detection to deliver auditory stimulation triggered by spontaneous activity in rodent electrophysiological signals using the software ‘Synapse’ (Tucker-Davis Technologies Inc., Alachua, FL, USA) (Fig.S6). The delivery of auditory stimuli was timed by the real-time detection of putative spindle events detected in one LFP signal per mouse recorded from the S1. Specifically, for each mouse, the recording channel showing highest incidence of spindles (layer 4) was chosen for real-time detection. The selected LFP signals were first filtered with a high-pass filter at 0.1Hz and low-pass filter at 100Hz and a second order parametric filter with centre frequency at 12.5Hz and a fractional bandwidth of 0.4 (octaves) was then applied (to filter the signal between 10-15Hz). Parametric filters are efficient for boosting the signal band of interest and making the attenuation of signals outside the selected band sharper, so their roll off (i.e., in our case: 9Hz or 16Hz) is low ^177^. We then calculated the square of the filtered signal and used an exponential smoothing function, which applies an exponentially decreasing weight to the data as a function of time. A threshold was then set to detect spindles based on the square of the filtered signal. The threshold for real-time detection was set at 4.5 times the mean of the smoothed power signal in line with previous automated detection algorithms^43, 44, 174^.

In order to restrict the detection of spindles to NREM sleep (i.e. avoiding REM sleep and movement), we set two conditions that had to be met for the algorithm to detect a putative ‘spindle event’. First, we calculated the square of the EMG signal, and a threshold was set for each mouse to distinguish between movement and immobility. Second, we applied a second order parametric filter to the occipital EEG channel, to filter the signal between 4-7Hz (i.e. theta frequency range, which is prominent during REM sleep^178–180^ and calculated online the square of this filtered signal. A corresponding threshold was set on this filtered signal for each mouse, to identify REM sleep. If either of these two conditions were met (i.e. mobility or high theta power in the occipital derivation) no putative ‘spindle event’ was detected.

#### Auditory stimulation

Open-field auditory stimulation was performed in the home cages, where mice were single-housed. The home cages consisted of 390 x 410 x 350mm electromagnetic shielded and sound-attenuated Faraday chambers (Lafayette Instrument, US). Sounds were played through magnetic speakers (MF1 Multi-Field Magnetic Speakers, Tucker-Davis Technologies) mounted on the chamber ceilings. Auditory stimuli were designed and triggered with the software Synapse (Tucker-Davis Technologies Inc., Alachua, FL, USA).

A pilot auditory-stimulation session during sleep was performed in three mice to identify sound parameters that generated an evoked response in S1 without inducing a state of arousal. Fifty different sounds, which ranged in frequency (between 4kHz–16kHz) and intensity (between 60dB–90dB) were used for this pilot. Pure tones played at 12kHz, for 100ms with an intensity of 70dB reached the best compromise, and therefore these parameters were used for the real-time stimulation. Additionally, a sham stimulation condition of 12kHz pure tones played at 0dB for 100ms was used as control. The sound intensity was calibrated using a sound level meter and calibration kit (Grainger, US).

The sound and sham condition were presented on two different days and the order of presentation was counterbalanced across mice. Each day, auditory stimulation was performed for six hours during the light period, specifically between ZT3.5 and ZT9.5. Sounds were delivered during (‘spindle’ condition) or in the absence of spindles (‘no-spindle’ condition) in a pseudo-random order. A minimum inter-stimulus interval of 3-s was allowed. Approximately 550 ± 30 stimuli (n=6 mice) were played during the 6-h of auditory stimulation. Figure (Fig.7A) shows examples of ‘spindle’ and ‘no-spindle’ trials.

Manual sleep scoring and offline automated spindle detection based on autoregressive (AR) modelling^10, 14^ were used to evaluate the performance of the online spindle detection and stimulation algorithm. Overall, 98.3% ± 0.3 of the auditory stimuli were presented during NREM sleep. Additionally, the comparative sensitivity (comparative true-positive rate) between detections made with the real-time detector and offline detections with the lowest threshold of the AR-model (r*_b_*=0.92), reached 86.2% ± 2.11. We calculated the EMG response to auditory stimulation across conditions (real sound vs sham condition presented during [‘spindle condition’] or outside [‘no-spindle condition’] spindle events). For the spindle condition only auditory stimuli presented during the occurrence of spindle events confirmed by detection by the AR-model ^10, 14^ were included in the analysis.

### Statistical Analyses

Data were analysed using MATLAB and its Statistics Toolbox, (The MathWorks, Inc) and IBM SPSS Statistics (IBM Corp). Linear mixed models, analysis of variance (ANOVA), factorial repeated measures ANOVA, repeated measures ANOVA, t-tests, and respective non-parametric tests were used as appropriate. To assess differences between specific groups, post-hoc tests were performed. The Tukey test was used to compare between groups when equal variances were assumed. The Sidak test was used to do multiple comparisons in cases where equal variances were assumed. Finally, the Games Howell test was used when equal variances were not assumed.

The parametric analyses mentioned above (ANOVA-based, t-test, and linear mixed models) require dependent variables and residuals to be normally distributed (although they are robust to violations in this assumption when group sizes are equal)^181^. Shapiro-Wilk normality tests were used to determine whether the data were normally distributed. In cases where the normality assumption was highly violated (and the size of the compared groups was different) either the data was transformed or non-parametric statistics were performed (i.e. Friedman test or Games-Howell post hoc test).

Mixed and multivariate tests require the variances for each combination of the groups to be homogenous. The Levene statistic was used to test for homogeneity of variance in the different assessed variables^181^. In cases where the homogeneity of variance assumption was violated, Welch’s F test or non-parametric posthoc tests (Games-Howell) were used. In the case of mixed models and repeated measures, the variances of the differences between groups of the within-subject factor (across the between-subjects factor) is required to be homogenous (i.e. sphericity assumption)^181^. Mauchley’s test of sphericity was used to assess whether the population variances of all possible different variable combinations were equal. In cases where the sphericity assumption was violated, the Greenhouse-Geisser (referred to as ‘GG’) correction was applied. This method corrects for the inflation in the F-value caused by lack of sphericity (unequal population variance at all variable levels) by multiplying the Greenhouse– Geisser estimate by the degrees of freedom used to calculate the F-value ^181^. In tables and figures, significance levels are indicated with black asterisks as follows: *p<0.05, **p<0.01, ***p<0.001.

---

**Supplementary Fig. S1.**
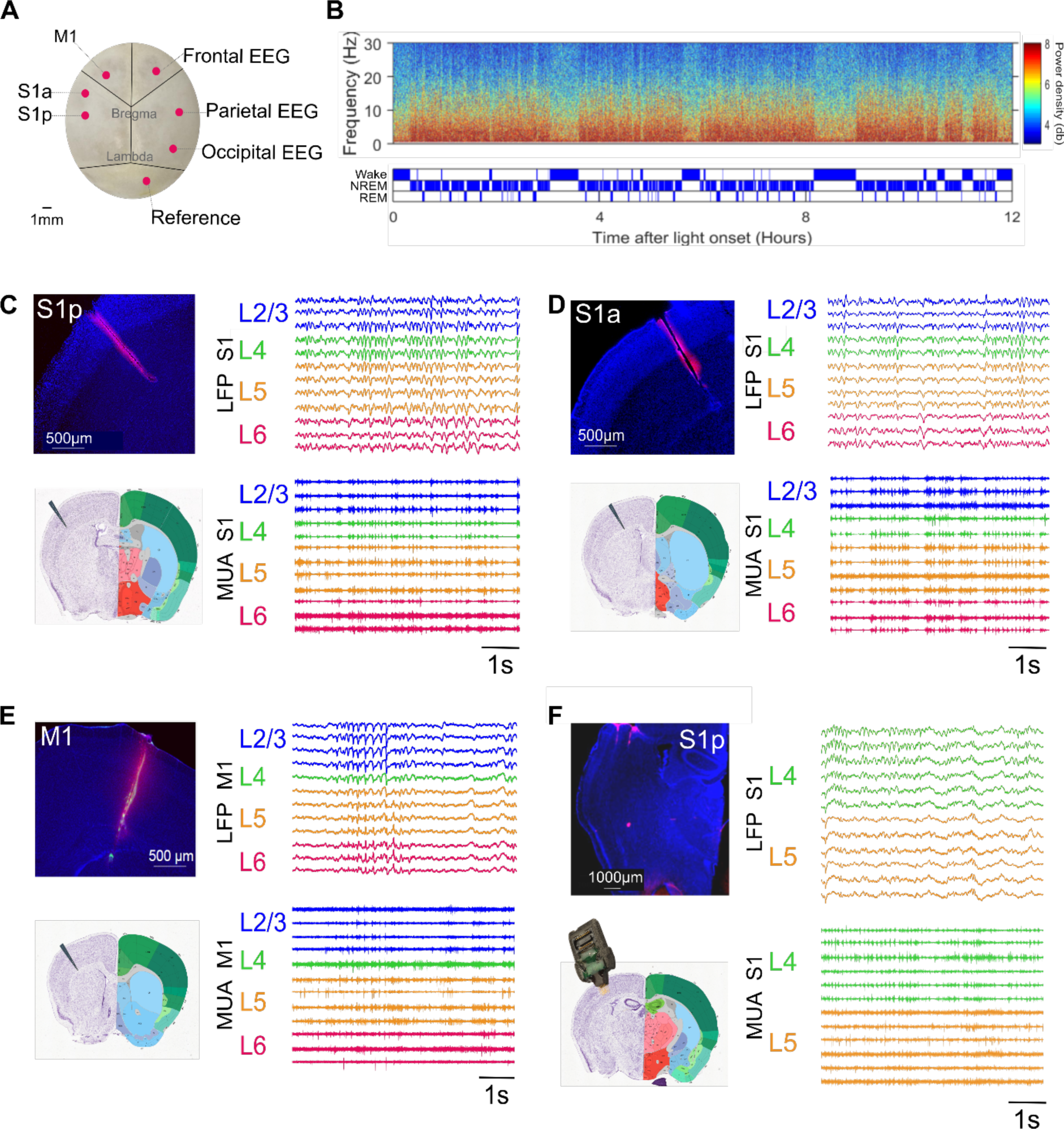
Recording sites and recordings of cortical activity during wake and sleep. **(A)** Locations where the EEG screws (frontal, parietal or occipital), LFP laminar probe or micro-wire arrays (M1, S1 anterior, S1 posterior) and reference screw (cerebellum) were implanted. **(B)** Spectrogram (top) and respective hypnogram (bottom) for one mouse during an undisturbed 12-hour light period. The spectrogram is colour-coded on a logarithmic scale. **(C,D,E,F)** Twenty-one mice were implanted with either: a 16-channel laminar probe in S1 anterior cortex (n=7) (C), S1 posterior cortex (n=7) (D), M1 cortex (n=7) (E), or a 16-channel micro-wire array in deep layers of S1 (n=7) (F). In (C,D,E,F), the location of the electrodes across or within S1 and M1 cortical layers was verified with histology of electrolytic microlesions and Dil stain electrode traces (magenta) on brain slices stained with DAPI (blue). The recording coordinates for each implant type were identified using the Allen mouse brain atlas. Representative 7-s segments of LFP and MUA signals (recorded during NREM) sleep across electrodes is shown for the different implants. (*Note:* EEG: electroencephalogram. LFP: local field potential. M1: primary motor cortex. S1: primary sensory cortex. MUA: multi-unit activity).

**Supplementary Fig. S2.**
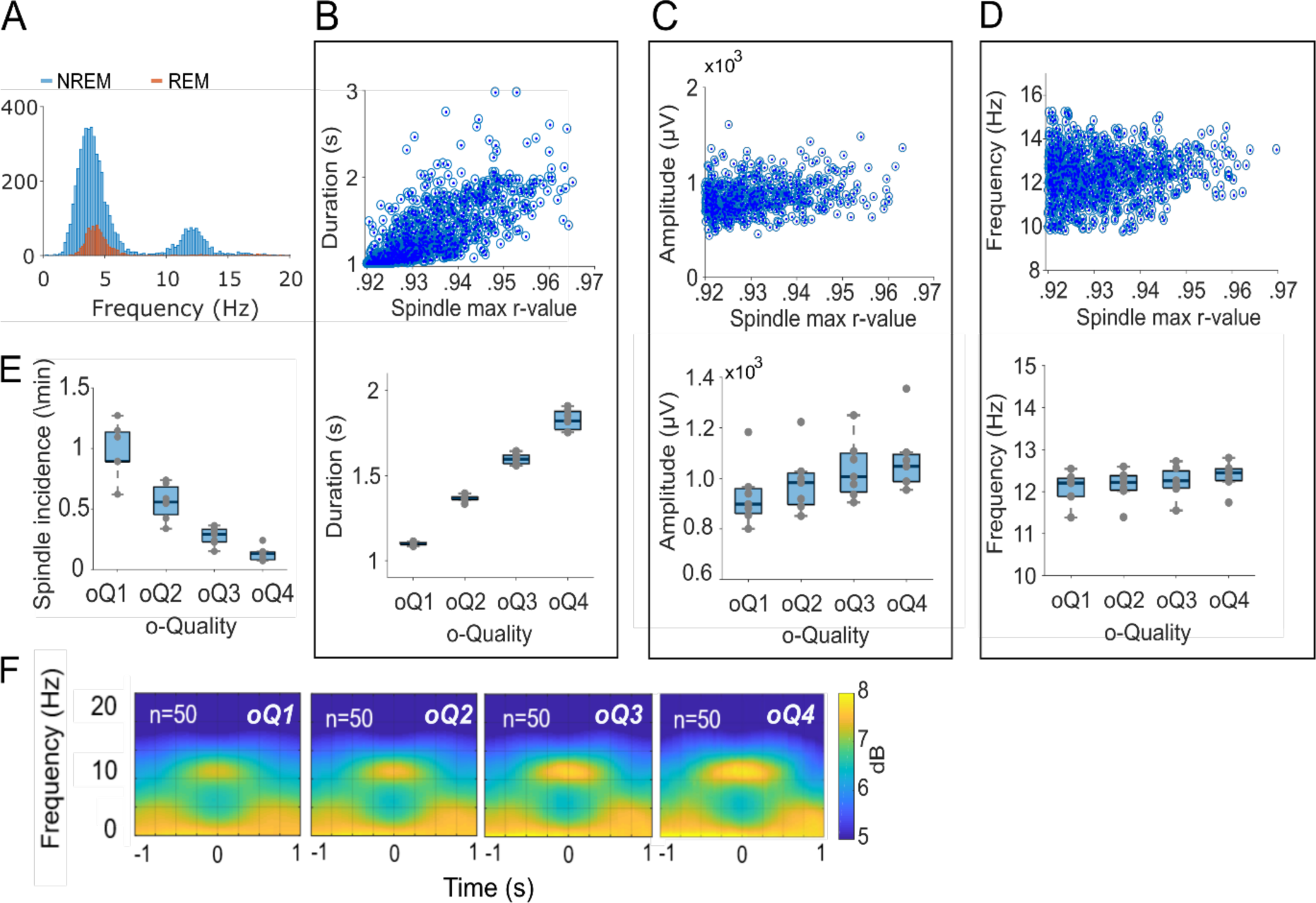
Properties of spindles as a function of their *o-Quality.* **(A)** Frequency distribution of all the oscillatory events detected by the AR-model during NREM (blue) and REM (red) sleep from electrodes located in layer 4 of S1 (the cortical layer with highest spindle density). **(B-D)** *Top:* Representative examples of the distribution of spindle duration, amplitude, and frequency as a function of the maximum *r*-value for each detected spindle in one mouse. *Bottom:* Mean duration, amplitude, and frequency of spindles with different *o-Quality*. **(E)** Spindle incidence per minute as a function of their *oQuality.* **(F)** Average spectrograms of 2-s segments of LFP signals recorded from electrodes located in layer 4 of S1. Spectrograms are aligned to the midpoint of each spindle event and averaged across 50 random spindles per o-Quality level and across mice. (Note: For figures B-D: n=7 mice. For boxplots: black lines= mean across mice, boxes= SEM, whiskers= 95% confidence intervals, dots= individual values for each mouse. AR: autoregressive. LFP: local field potential. S1: primary somatosensory cortex. SEM: standard error of the mean).

**Supplementary Fig. S3.**
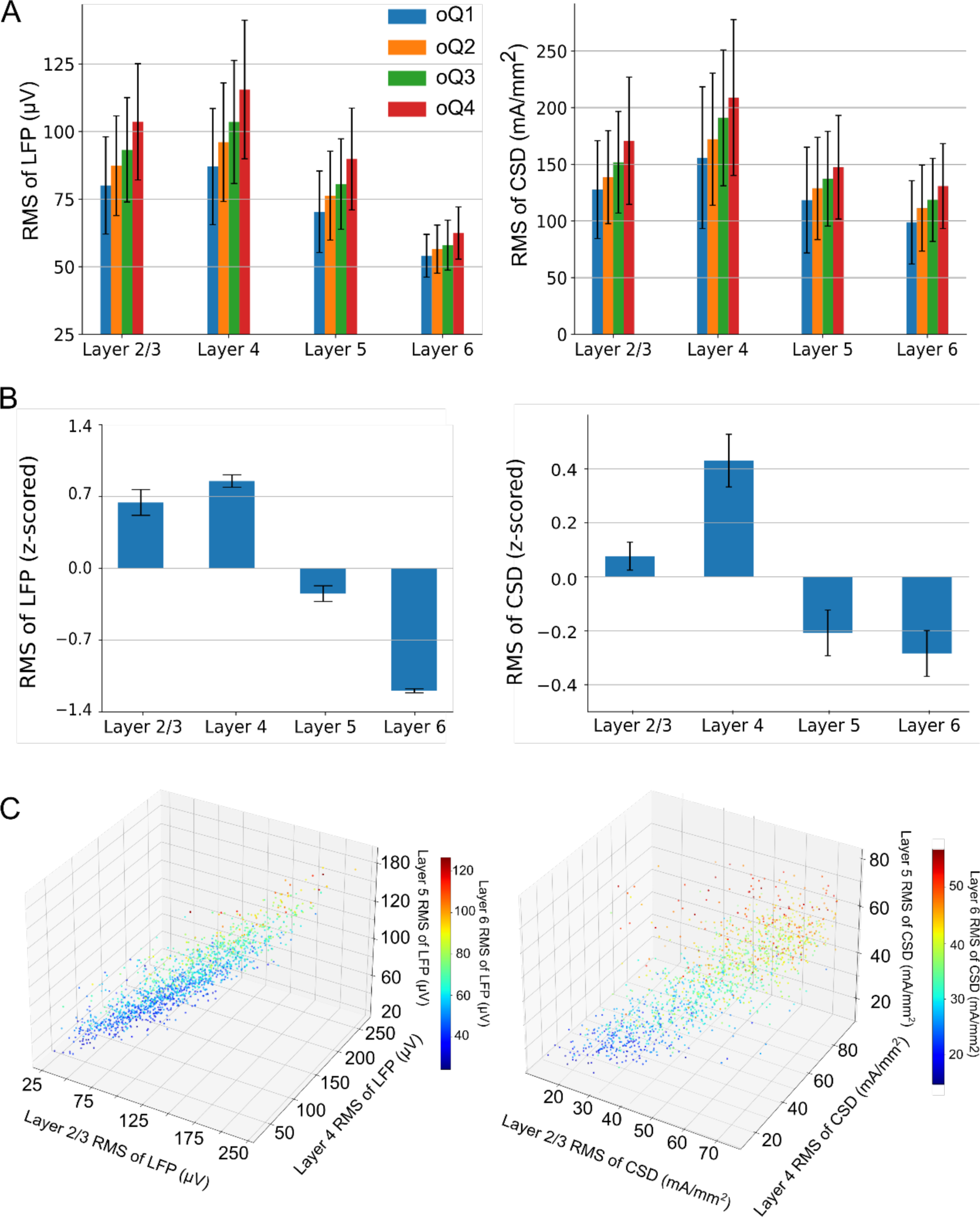
RMS and CSD profiles of spindles across layers. **(A)** Laminar profile of LFP (left) and CSD (right) in an example mouse, for each spindle quality. The laminar profile is represented by the average 1-s root mean square (RMS) value from the spindle centre, averaged across channels within each layer. Error bars represent the standard deviation across spindles. **(B)** LFP (left) and CSD (right) laminar profile of all spindles, grand average across mice (n=7). For each mouse, RMS values are z-scored across all cortical channels, and subsequently averaged across channels within each layer and across spindles. Error bars indicate the standard error of the mean. **(C)** 3-D scatterplots showing LFP (left) and CSD (right) RMS for each layer in an example mouse. Each point represents one spindle, with layers 2/3, 4 and 5 plotted onto the x-axis, y-axis and z-axis respectively, and layer 6 represented by the colour scale. Strong collinearity across layers can be observed for both LFP and CSD. (*Note:* LFP: local field potential. CSD: current source density).

**Supplementary Fig. S4.**
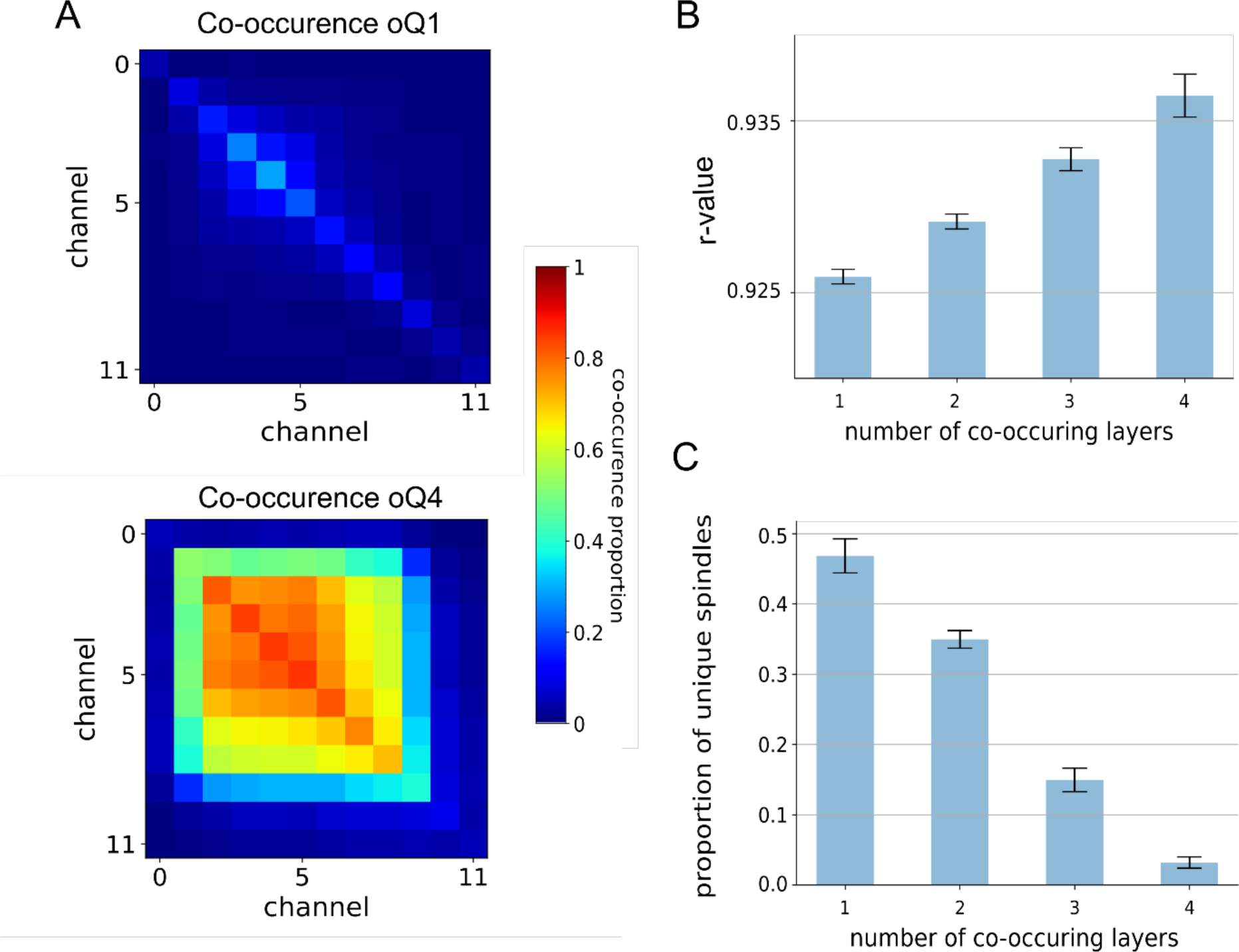
Co-occurrence of spindles across cortical layers. **(A)** Co-occurrence matrix in an example mouse for spindles of quality one (top) and quality four (bottom). Colour scale indicates the proportion of spindles from each quality that were detected in each channel pair simultaneously. **(B)** Mean quality metric of spindles (r-values) co-occurring in different layers simultaneously in one example mouse. Error bars indicate the standard deviation. **(C)** Group mean across mice of proportion of spindles co-occurring in different layers (n=7). Error bars indicate standard error of the mean (SEM).

**Supplementary Fig. S5.**
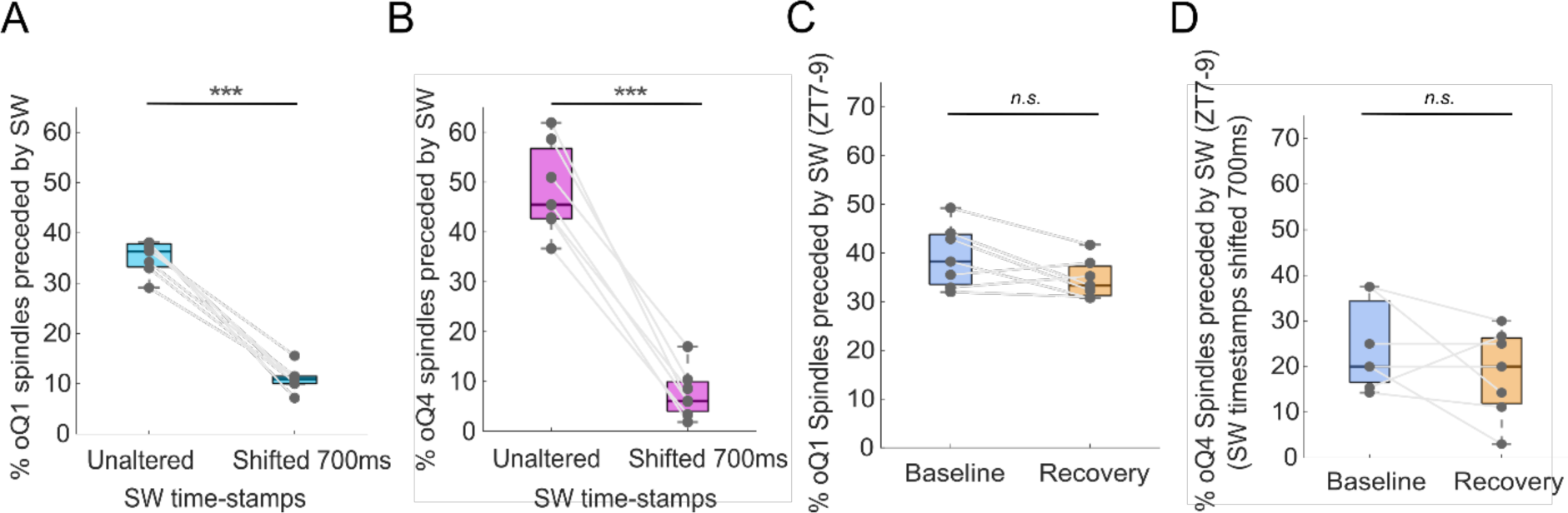
Coupling of spindles and SW after shifting the timestamps of SW. **(A)** Percent of low *o-Quality* (oQ1) spindles preceded by SW when SW timestamps (i.e. the start time of automatically detected individual depth-positive high-amplitude SW, see methods ‘*Slow wave detection*’) were unaltered or shifted forward by 700ms. **(B)** Percent of high *o-Quality* spindles (oQ4) preceded by SW when SW time-stamps were unaltered or shifted by 700ms. **(C)** Percent of low *o-Quality* (oQ1) spindles preceded by SW during the first two hours of sleep recovery (ZT7-ZT9) after 6 hours of sleep deprivation (orange) and corresponding baseline sleep time-period (blue). **(D)** Percent of high *o-Quality* (oQ4) spindles preceded by SW (with timestamps shifted by 700ms) during the first two hours of sleep recovery (ZT7-ZT9) after 6 hours of sleep deprivation (orange) and corresponding baseline sleep time-period (blue). (*Note:* For boxplots: black lines= mean across mice, boxes= SEM, whiskers= 95% confidence intervals, dots= individual values for each mouse. SW: slow waves. ZT: zeitgeber time. SEM: standard error of the mean).

**Supplementary Fig. S6.**
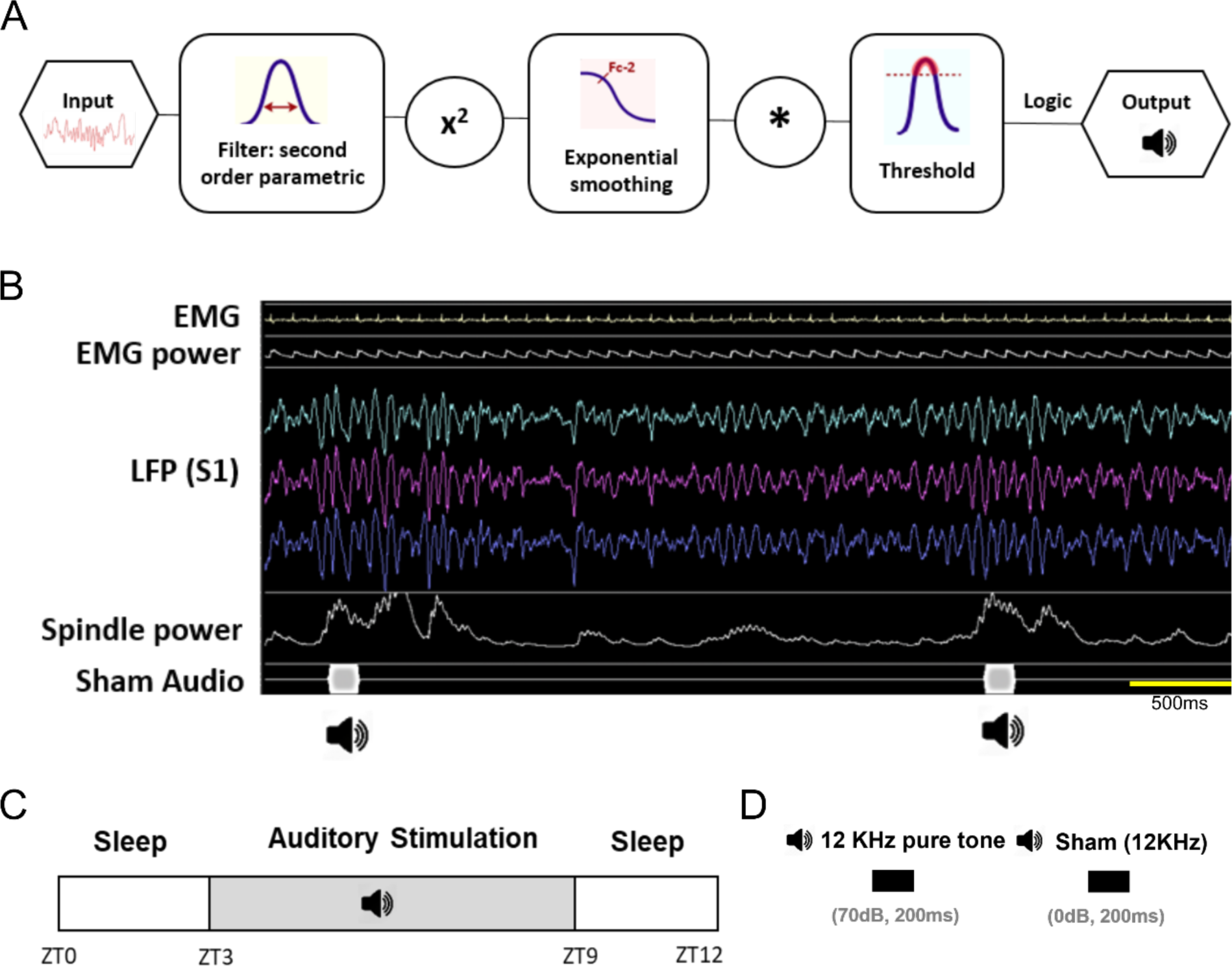
Principles and schematic of the auditory closed-loop stimulation paradigm. **(A)** Processing steps followed by the system to detect spindles (10-15Hz) in real time and deliver sounds times-based on these detections. **(B)** Examples of real-time spindle detection in a sham (0dB sound) condition. Spindles were detected when the power of S1 LFP signals increased and reached a predefined threshold. Sounds were delivered only if the EMG and theta power in the occipital EEG were low. **(C)** Schematic of the structure of the auditory-stimulation paradigm performed using the closed-loop system, across 12h light periods (white bar). Zeitgeber Time (ZT) indicates the time from light onset (lights on at 9am; lights off at 9pm). The shaded boxes indicate the times during which the auditory stimulation paradigms were carried out. **(D)** Auditory stimuli consisted of 12kHz pure tones presented at 70 dB or 0 dB (sham) for 200ms. (*Note:* S1: primary somatosensory cortex. LFP: local field potential. EMG: electromyography).

**Supplementary Fig. S7.**
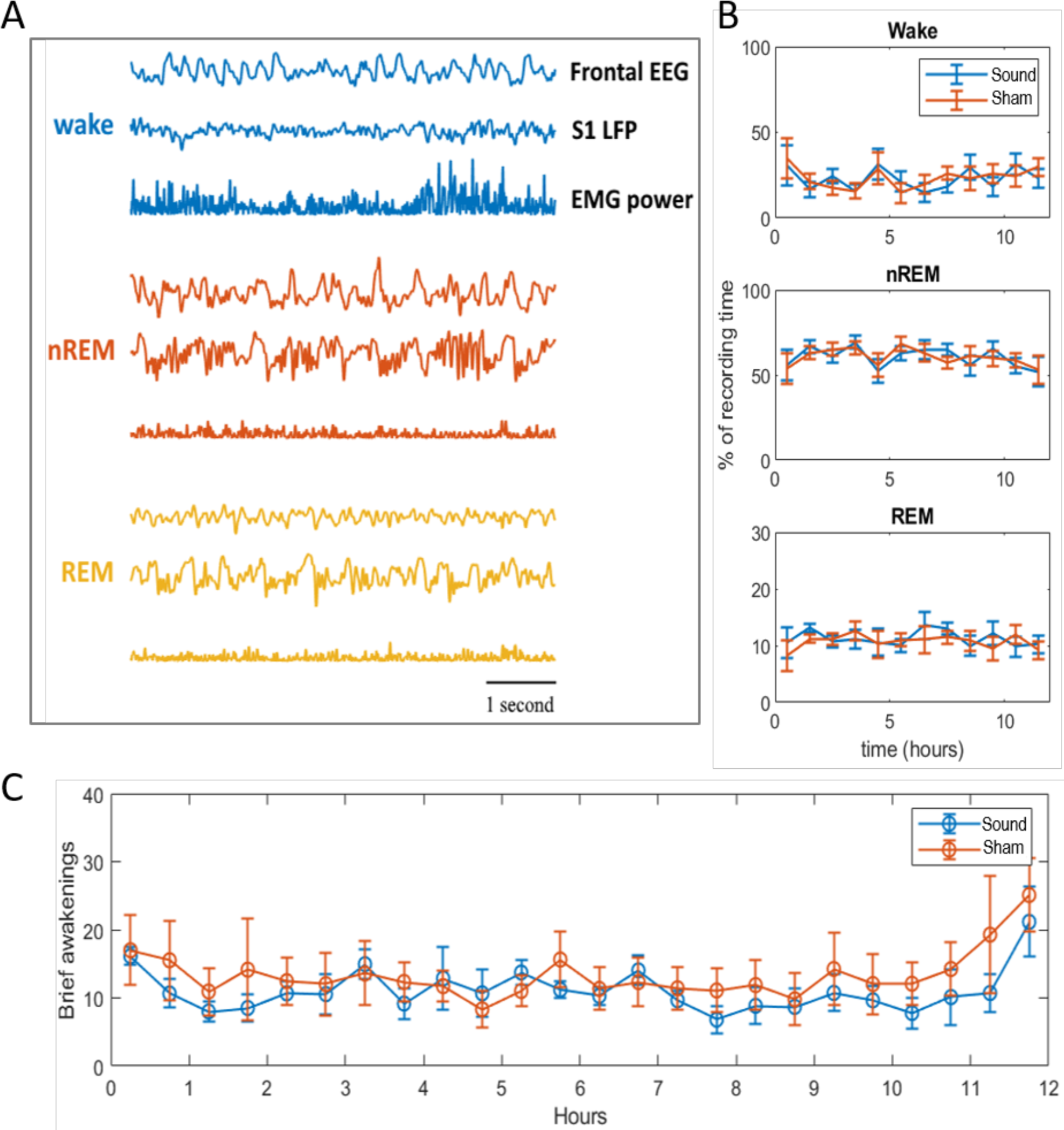
Auditory stimulation does not affect vigilance states. **(A)** Representative frontal EEG, S1 LFP, and EMG power traces during wake, NREM, and REM sleep in one mouse. **(B)** Time course of vigilance states over the 12h recording period, in 1h intervals for real stimulation and mock. The amount of each state is represented as a percentage of the total recording time. Mean ± SEM (n=6 mice). **(C)** Brief awakenings during the 12h recording period, shown as number/hr of NREM sleep. (*Note*: EEG: electroencephalography. S1: primary somatosensory cortex. LFP: local field potential. EMG: electromyography. SEM: standard error of the mean).

**Supplementary Fig. S8.**
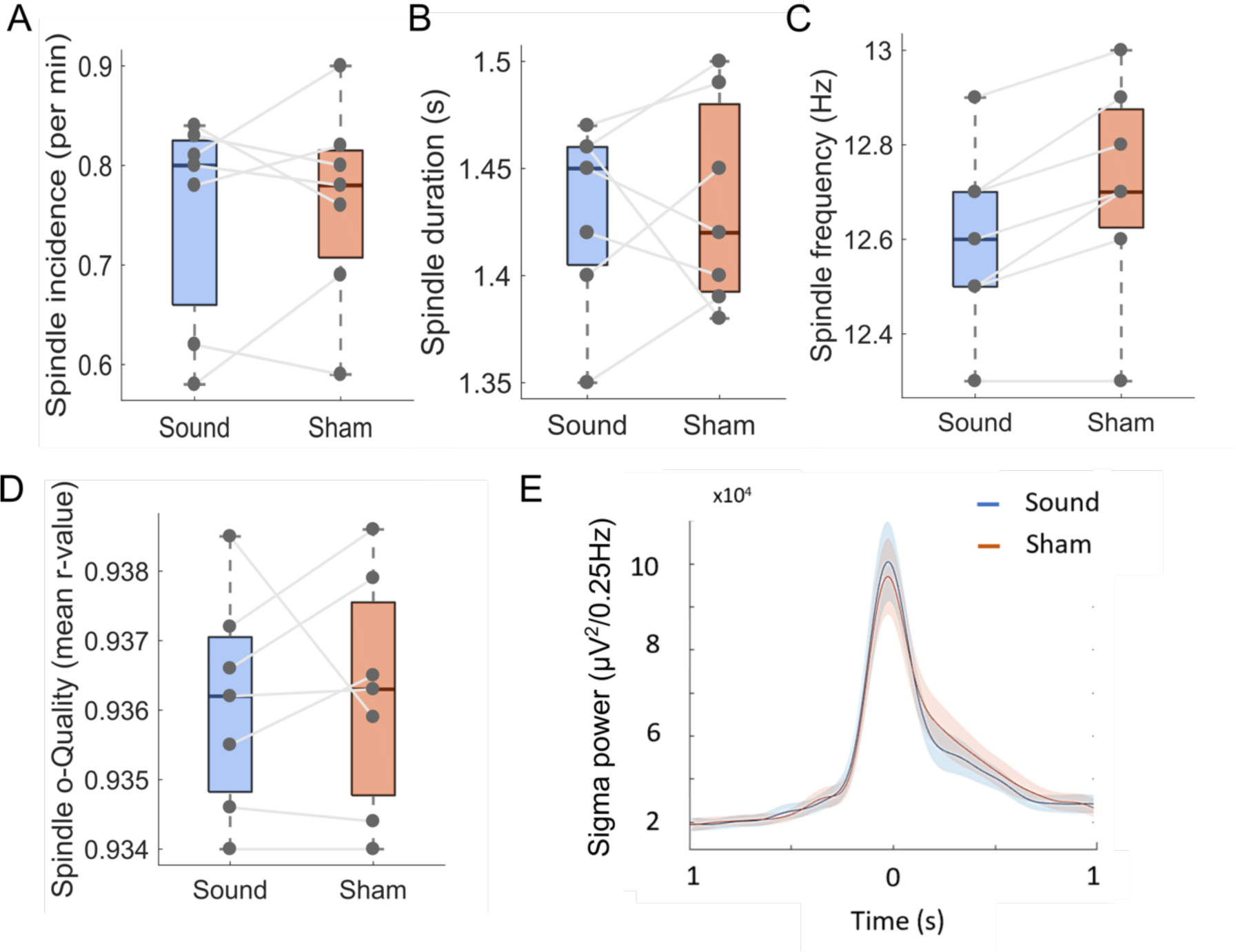
Spindle metrics are not affected by pure tones. Mean spindle incidence **(A),** duration **(B),** frequency **(C),** and *o-Quality* (higher r-value = higher *o-Quality*) **(D)**, across all mice (n=7) for spindles coincident with auditory stimulation and spindles coincident with sham stimulation. **(E)** Mean ± SEM sigma power time course, where sound stimulation or sham stimulation occurs at time 0s. (*Note:* Lines= average across mice, shaded area=SEM. (*Note:* SEM: standard error of the mean).

**Supplementary Fig. S9.**
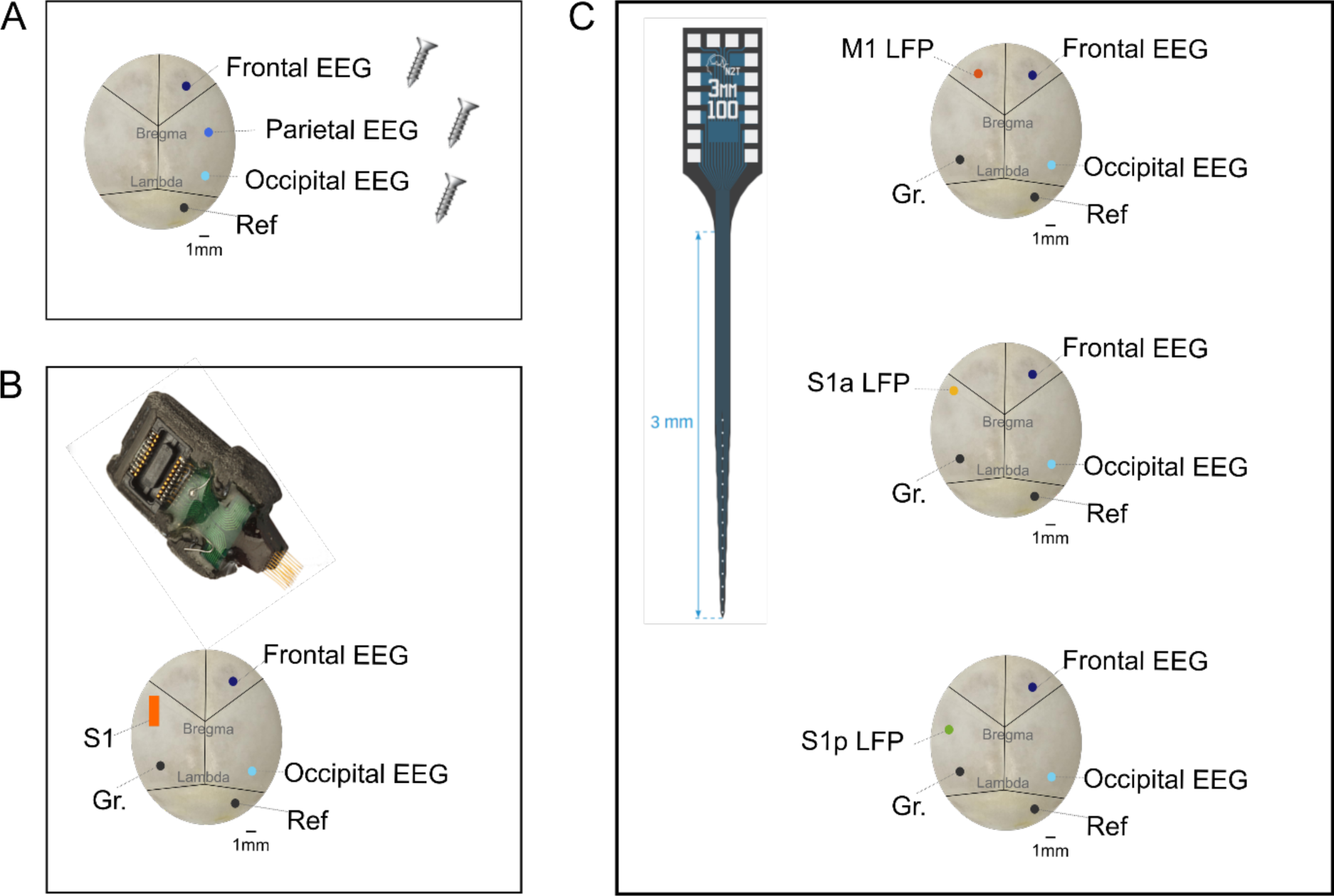
Electrode placement configurations. **(A-C)** Locations where the EEG screws, LFP laminar probe or LFP micro-wire array, reference screw (Ref) and ground screw (Gr) were implanted. **(A)** A total of n=6 C57Bl/6 mice were implanted with EEG screws epidurally above the frontal, parietal and occipital cortices. **(B)** A total of n=21 mice (n=7 C57/BL6; n=7 GRIA1-/-; n=7 WT littermates) were implanted with frontal and occipital EEG screws plus a polyimide-insulated tungsten microwire array into deep layers of S1 (layers 4-5). **(C)** A total of n=21 C57Bl/6 mice were implanted with frontal and occipital EEG screws plus a 16-channel laminar probe into the primary motor cortex (M1, n=7), the anterior part of the primary somatosensory cortex (S1a, n=7) and the posterior part of the primary somatosensory cortex (S1p, n=7). (*Note:* EEG: electroencephalography. LFP: local field potential. WT: wild-type. M1: primary motor cortex. S1: primary somatosensory cortex.).

**Supplementary Fig. S10.**
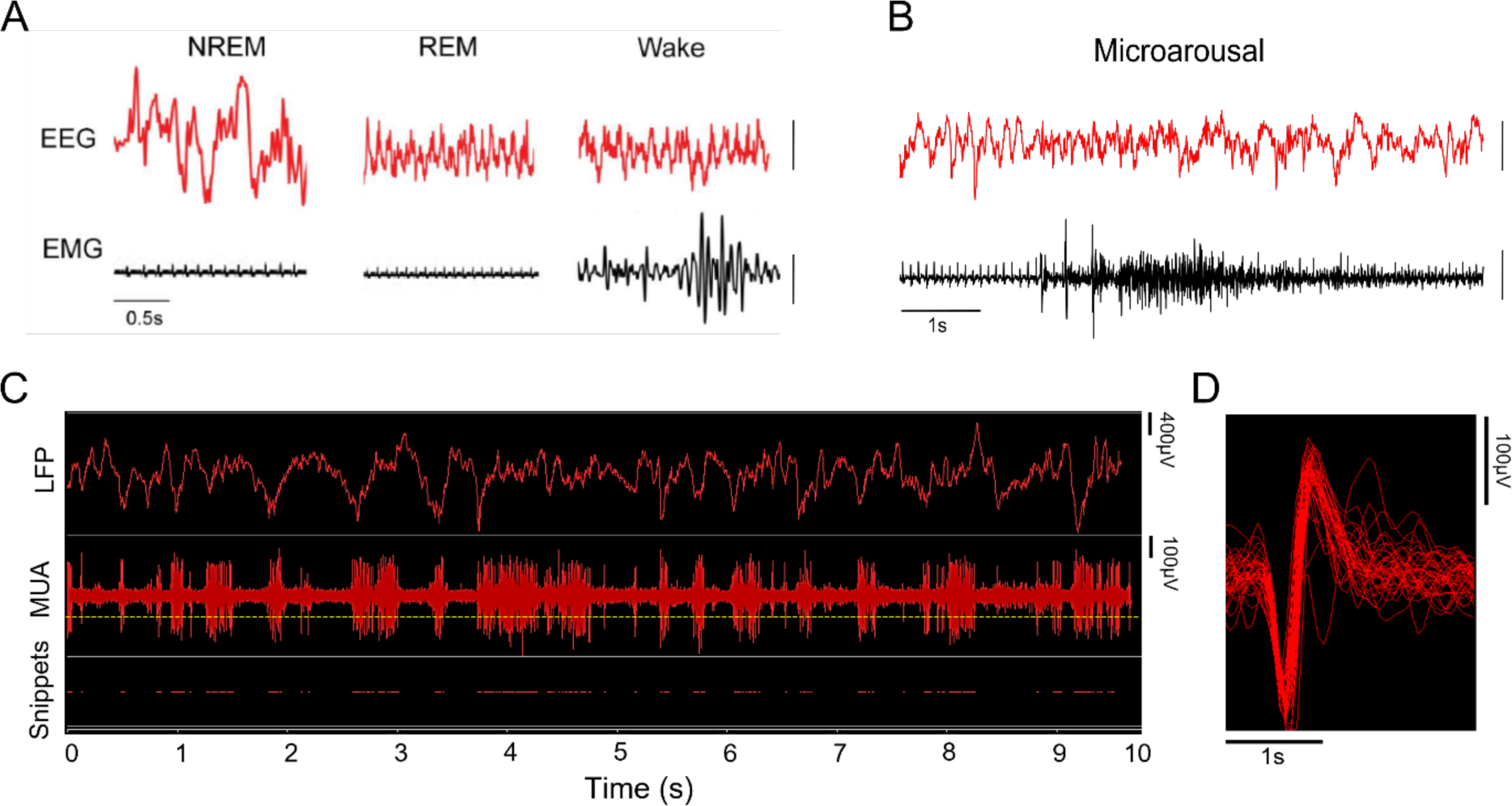
Sleep stages, multi-unit activity and spike waveform. **(A)** Representative EEG signal segments from the frontal cortex (top) and the EMG (bottom) recorded in one mouse during different vigilance states (scale bar=250µV). (**B)** Representative EEG (frontal) and EMG traces recorded from one mouse during a microarousal. Microarousals were defined as transient periods of low voltage, high frequency oscillations in the EEG signals accompanied by elevated EMG tone, lasting ≥4s and ≤16s. **(C)** Representative LFP (top), MUA (middle) and snippets (timestamped spike waveforms) recorded simultaneously from S1 in one mouse during NREM sleep. The yellow dotted line represents the amplitude threshold used for spike acquisition. When the recorded voltage of the MUA crossed this threshold, 46 samples around the event (0.48 ms before, 1.36 ms after the threshold crossing) were extracted. **(D)** Corresponding waveforms of the action potentials recorded extracellularly. (*Note:* EEG: electroencephalography. EMG: electromyography. LFP: local field potential. MUA: multi-unit activity).

**Supplementary Fig. S11.**
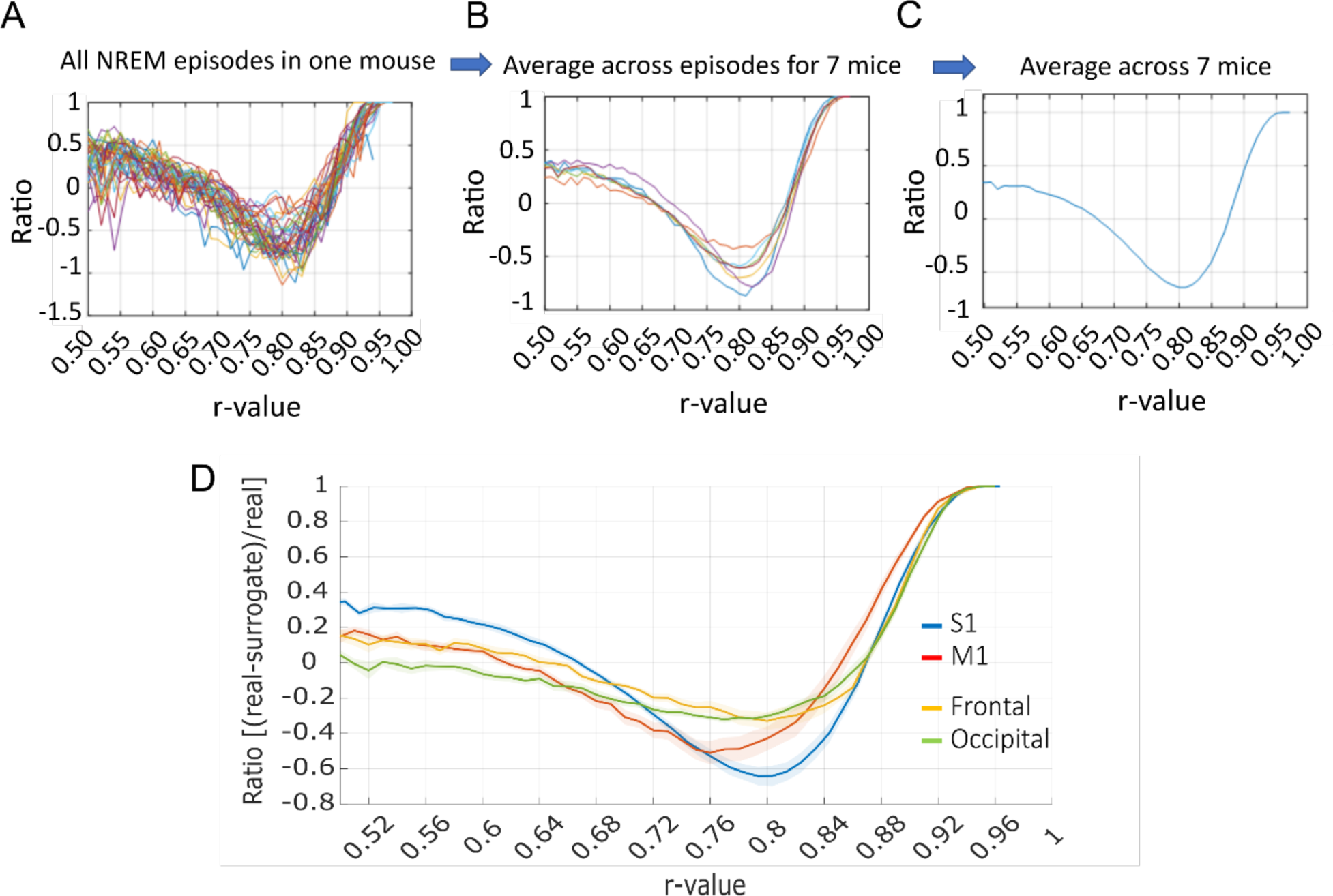
Validation of the upper-threshold (rb). **(A)** The continuous distribution of r-values (negative logarithm proportional to damping constant) for a pole with frequency of 10-15Hz was calculated for every NREM episode and its corresponding surrogate signal. Here we show the rate (i.e. real signal – surrogate signal /real signal) between the r-value distributions in the real signals and their corresponding surrogates for all NREM episodes detected in one example mouse in the S1 LFP signal. Each colour represents an individual NREM episode. **(B)** Same as *A* but averaged across NREM episodes for n=7 mice. Each colour represents an individual mouse. **(C)** Same as *B* but averaged across mice. **(D)** Ratio between the r-value distributions in NREM episodes of real signals and respective surrogates recorded from intracortical channels (S1 and M1) and the EEG (frontal and occipital). Lines= average across NREM episodes and mice (n=7 per derivation), shaded area=SEM. (*Note:* S1: primary somatosensory cortex. LFP: local field potential. M1: primary motor cortex. EEG: electroencephalography).

**Supplementary Fig. S12.**
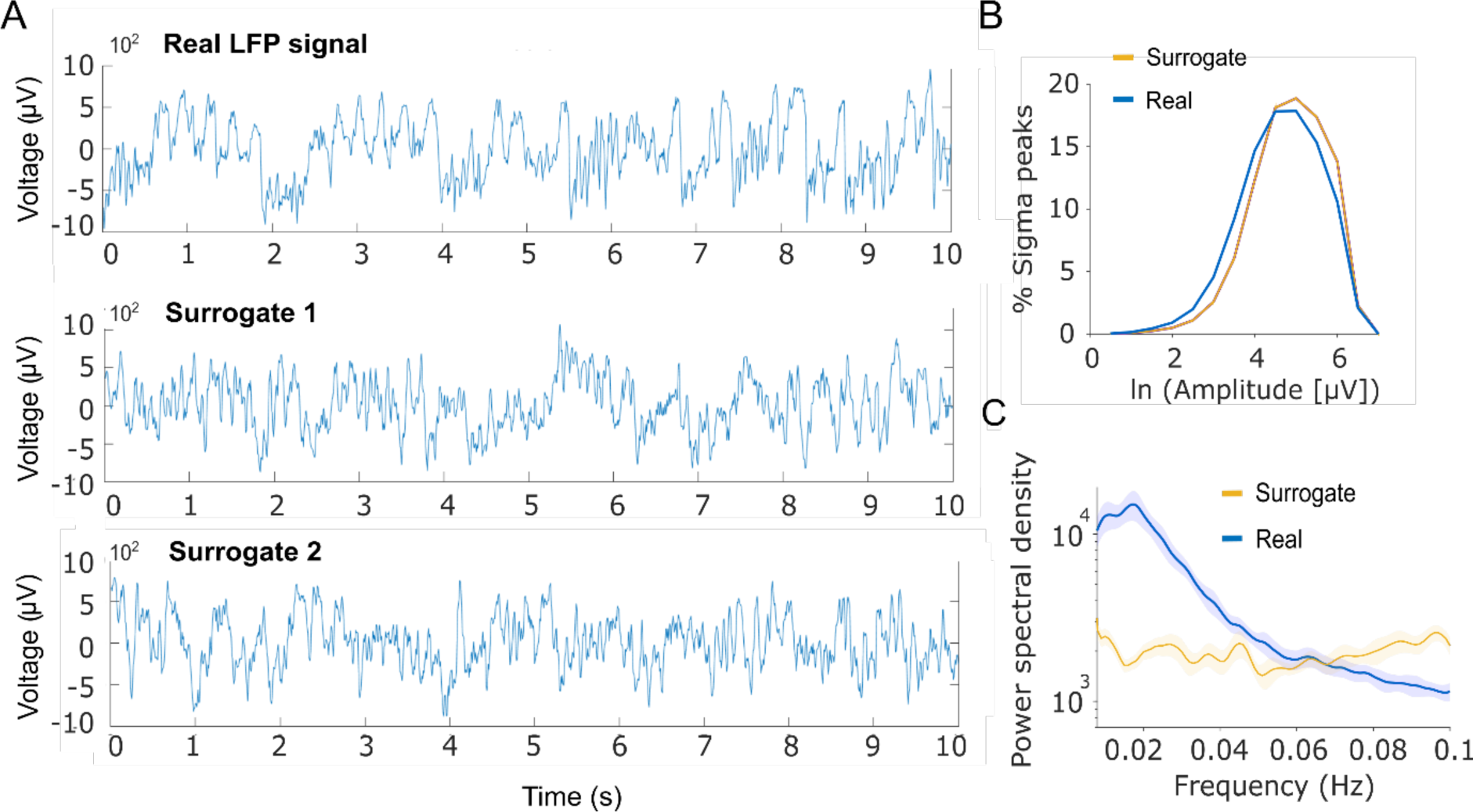
Examples and properties of surrogate signals. **(A)** Example timeseries of a real LFP signal recorded from S1 in one mouse and two surrogate signals created based on an improved version of the IAAFT (Iterative Amplitude Adjusted Fourier Transform) algorithm. **(B)** Peak sigma (10-15 Hz) amplitude distribution calculated from a 10-minute segment of LFP real signal (in one mouse) and 19 respective surrogate signals. Shaded area= SEM. **(C)** Power spectral density (µV^2^/0.25Hz) of the envelope (Hilbert transform) of filtered LFP signals (10-15 Hz) recorded from S1 and respective filtered (10-15 Hz) surrogates (n=19 per derivation per mouse). Figure shows mean and SEM (shaded area) across mice (n=7). (*Note:* LFP: local field potential. S1: primary somatosensory cortex. SEM: standard error of the mean).

## Acknowledgements

We thank all members of the laboratory of VVV for their help with surgery assistance, animal care and sleep deprivation; Thomas Jahans-Price and Marios Panayi for their help managing the GRIA1 mouse colony and establishing the microlesion protocol; the TDT support team (Myles Billard, Mark Hanus, Victor Rush) for their technical support with electrophysiology data acquisition; Kristina Parley for her support with the histological procedures; Fernando Nodal for his help setting up the auditory equipment; David Dupret for his scientific advice. This work was supported by the Wellcome Trust PhD studentships 109059/Z/15/Z (CBD), 203971/Z/16/Z (LBK), the Medical Research Council (UK) grant MR/S01134X/1 (VVV) and the Wellcome Trust grants 106174/Z/14/Z and 098461/Z/12/Z. CBD was also supported by a Clarendon Scholarship (provided by the University of Oxford). ECPG was supported by a Winton Exoplanet Fellowship. MCK and JPD were supported by Berrow Foundation Lord Florey Scholarships.

## Author contributions

CBD, VVV and DMB proposed, designed, and initiated the study. CBD and LBK soldered EEG recording implants and cables. CBD, LBK and MCK performed mouse surgeries. CBD conducted the electrophysiology recordings and experiments on mice. CBD and LBK performed microlesions, perfusions and histology. DMB provided the transgenic mice. EO and PA developed the spindle detection algorithm. CBD, SAB, JPD analysed the data. VVV, EO, PA, ECPG, DMB and EOM guided the data analysis. CBD and VVV wrote the manuscript. All the authors discussed the results and commented on the manuscript.

## Competing interests

The authors declare no competing interests.

## Notes

### Competing Interest Statement

The authors have declared no competing interest.

